# Rhodoquinone-dependent electron transport chain is essential for *C. elegans* survival in hydrogen sulfide environments

**DOI:** 10.1101/2024.02.23.581771

**Authors:** Laura Romanelli-Cedrez, Franco vairoletti, Gustavo Salinas

## Abstract

Hydrogen sulfide (H2S) has traditionally been considered as an environmental toxin for animal lineages; yet, it plays a signaling role in various processes at low concentrations. Mechanisms controlling H2S in animals, especially in sulfide-rich environments, are not fully understood. The main detoxification pathway involves the conversion of H2S into less harmful forms, through a mitochondrial oxidation pathway. The first step of this pathway oxidizes sulfide and reduces ubiquinone (UQ) through sulfide-quinone oxidoreductase (SQRD/SQOR). Because H2S inhibits cytochrome oxidase and hence UQ regeneration, this pathway becomes compromised at high H2S concentrations. The free-living nematode *C. elegans* feeds on bacteria and can face high sulfide concentrations in its natural environment. This organism has an alternative ETC that uses rhodoquinone (RQ) as the lipidic electron transporter and fumarate as the final electron acceptor. In this study, we demonstrate that RQ is essential for survival in sulfide. RQ-less animals (*kynu-1* and *coq-2e* KO) cannot survive high H2S concentrations, while UQ-less animals (*clk-1* and *coq-2a* KO) exhibit recovery, even when provided with a UQ-deficient diet. Our findings highlight that *sqrd-1* uses both benzoquinones and that RQ-dependent ETC confers a key advantage (RQ regeneration) over UQ in sulfide conditions. *C. elegans* also faces cyanide, another cytochrome oxidase inhibitor, whose detoxification leads to H2S production, via *cysl-2*. Our study reveals that RQ delays killing by the HCN-producing bacteria *Pseudomonas aeruginosa* PAO1. These results underscore the fundamental role that RQ-dependent ETC serves as a biochemical adaptation to H2S environments, and to pathogenic bacteria producing cyanide and H2S toxins.

## Introduction

Hydrogen sulfide (H2S) has long be regarded as toxic to many animals due to inhibition of cellular respiration(1). Recent research in mammals has shown that H2S can function as a signaling molecule at low concentrations(2), renewing interest in H2S research. Other animal lineages, such as invertebrates that thrive in marine sediments, can cope with environmental hypoxia and sulfide-rich environments(3–6). The molecular mechanisms that underlie the control of H2S levels differ in different animal lineages and are not fully understood.

In animals, the main detoxification pathway of H2S involves its conversion into less toxic forms, through a mitochondrial oxidation pathway(7). The first enzyme involved in this pathway is sulfide:quinone oxidoreductase (SQRD/SQOR), which catalyzes the oxidation of H2S to less toxic low molecular weight persulfides and the reduction of ubiquinone (UQ). The persulfides are further oxidized to sulfate in subsequent reactions catalyzed by persulfide dioxygenase, sulfur transferase and sulfite oxidase(7–9). Because SQRD transfers electrons to the electron transport chain (ETC) and H2S inhibits ETC complex IV (cytochrome c oxidase), this pathway may be insufficient to withstand high H2S concentrations. Invertebrates living in sulfide-rich environment, such as mussels and nematodes, have multiple SQRD/SQOR genes and additional mechanisms to deal with H2S, including association with episymbiont and endosymbiont bacteria(5).

*C. elegans* inhabits the soil, where it feeds on bacteria and encounters various environmental challenges, including bacteria that produce sulfide and/or cyanide(10–12). Indeed, cyanide poisoning is a killing mechanism used by *C. elegans* pathogens, such as *Pseudomonas* spp(10). These toxins may have imposed selective pressures leading to specific biochemical adaptations. Indeed, the cysteine synthases homologs CYSL-1 and CYSL-2 conform a cyanide-assimilation pathway. CYSL-2 is a cyanoalanine synthase that catalyzes the conversion of cyanide and cysteine into β-cyanoalanine and H2S, lowering cellular cyanide levels(13). The resultant H2S is detoxified mainly by the sulfide oxidation pathway. CYSL-1, in turn, is an enzyme that in some lineages catalyzes the conversion of H2S and O-acetylserine to cysteine and acetate, but in *C. elegans* functions as a sulfide sensor(11, 14).

In the presence of H2S, CYSL-1 interacts with the O2-sensing hydroxylase EGL-9(14, 15) to promote a HIF-induced response to H2S and HCN, which leads to an increased expression of *sqrd-1 and cysl-2*(16, 17). This suggests that this hypoxia-independent HIF-1 response in *C. elegans* evolved to withstand high concentrations of naturally occurring H2S and HCN. It has been proposed that *cysl-1* and *cysl-2* genes would have been co-opted to protect this lineage against cyanogenic toxins(13). More recently, a HIF-1-independent sulfide detoxification mechanism involving the SKN-1/NRF-2 transcription factor has been described, through CYSL-1 and RHY-1(18).

A key biochemical adaptation of *C. elegans* and some other animal lineages that face hypoxia is the existence of an alternative mitochondrial ETC in which rhodoquinone (RQ), and not UQ, serves as the lipidic electron transporter(19, 20). In this alternative ETC, complex I reduces rhodoquinone (RQ) to rhodoquinol (RQH2). RQH2 is oxidized back to RQ by complex II, which reduces fumarate to succinate, catalyzing the reverse reaction to the canonical activity of complex II (succinate dehydrogenase)(21, 22). In this abbreviated alternative ETC, fumarate, not oxygen, is the final electron acceptor(22). Because both H2S and HCN inhibit complex IV, this alternative ETC would confer an advantage to *C. elegans* survival against these toxins. *C. elegans* mutants unable to synthesize RQ are more sensitive to cyanide than mutants unable to synthesize UQ(23).

The structural difference between RQ and UQ is a single substituent at position 2 of the benzoquinone ring (an amine group in RQ vs a methoxy group in UQ), This change impacts in the standard reduction potential. The É° of RQ/RQH2 couple (rhodoquinone/rhodoquinol) has been reported to be -30 or -63 mV (bound to chromatophores or pure in a hydroalcoholic solution, respectively)(24). Because RQ/RQH2 É° is higher than that of NAD^+^/NADH couple (-320 mV) and lower than that of fumarate/succinate couple (+30 mV), it allows fumarate to be readily reduced by RQH2, and RQ by NADH. In contrast, the É° of UQ/UQH2 couple (ubiquinone/ubiquinol) is +50 mV or +43 mV (bound to chromatophores or pure, respectively)(24), higher than that of the succinate/fumarate couple, thus thermodynamically favoring succinate oxidation. Importantly, RQ is synthesized by nematodes, bivalves and annelids, which are the animal lineages more abundant in hypoxia and sulfide-rich environments(3–6).

We reasoned that the canonical sulfide detoxification pathway, which oxidizes sulfide and reduces ubiquinone would not be functional at high sulfide concentrations due to inhibition of complex IV, while the RQ-dependent ETC would be fully functional in sulfide conditions. Although UQ-complex II in reverse has been implicated in sulfide oxidation *in vitro*(25), the redox potential of RQ-complex II allows sulfide oxidation over a broader range of concentrations. In the present work, we demonstrate that RQ is essential for survival at high sulfide concentrations. Mutants unable to synthesize RQ are more susceptible to H2S than mutants unable to synthesize UQ. In addition, we showed that besides sulfide:quinone oxidoreductase *sqrd-1*, previously reported to be essential in *C. elegans* sulfide response, Y9C9A.16 (from now on *sqrd-2*), thought to be a pseudogen, is expressed and plays a marginal role in the recovery response to sulfide. Finally, we show that RQ has a role in the presence of the HCN-producing bacteria *Pseudomonas aeruginosa* PAO1.

## Results

Because sulfide at high concentrations inhibits complex IV of the canonical ETC we hypothesize that RQ would be the key benzoquinone in sulfide detoxification, accepting electrons from H2S through SQRD/SQOR. This would confer the advantage of transferring electrons independently of complex IV. SQRD/SQOR uses FAD as a cofactor to transfer electrons from H2S to UQ in the canonical mechanism. Because the standard reduction potential of FAD attached to the SQRD/SQOR is approximately -123 mV(43); it would be thermodynamically possible to transfer electrons from FAD to RQ, but whether the active site of *C. elegans* SQRD/SQORs can accommodate RQ is not known.

### C. elegans possess two sulfide:quinone oxidereductases that accommodate both RQ and UQ

*C. elegans* possesses two SQRD/SQOR genes, both with approximately 45 % identity to human SQOR (alignment shown in **Figure S1**). One of them, *sqrd-1*, has been previously characterized and involved in sulfide response(11), while the other, *Y9C9A.16*, was initially proposed to be a pseudogen (quoted in (11)). Interestingly, other nematode, platyhelminth and mollusk lineages also possess two SQRD genes (**Figure S1**). We found that *Y9C9A.16* is expressed in *C. elegans*, yielding an mRNA of the expected size according to the gene model (**Figure S2**).

We then examined in silico whether *C. elegans* SQRD-1 and SQRD-2 would be able to accommodate RQ and/or UQ at the benzoquinone-binding site. Both quinones bind similarly to *C. elegans* SQRD-1 and SQRD-2. There are no clashes that would impede the binding of either benzoquinones to either enzyme. Binding free energy estimated through GBVI and KDeep are negative and suggest a favorable interaction (**Table 1**). Both quinones establish hydrogen bonds through the carbonyl moieties with Trp455 and Lys438 (numbering corresponding to SQRD-1 isoform a, highlighted in cyan in **Figure S1**) in both enzyme models, being the main polar interactions with the enzyme (**Figure 1**, **A** and **B**). All the other interactions are hydrophobic and thus little affected by the presence of a methoxy or amino substituent. Furthermore, both quinones are located near the FAD cofactor in the holoenzyme, within a distance compatible with electron transfer reactions and within the range of quinone/FAD distance in PDB deposited structures (**Figure 1C).** The stability of the modeled SQRD complexes with RQ and UQ was tested using equilibrium molecular dynamics. The simulations showed that all four models equilibrated and remained stable at least for 50 ns. The secondary structure of the homology models was kept during the simulation and the complexes evolved similarly when compared to the crystal structure 6oic (also dynamized under the same conditions). UQ and RQ were kept in the quinone site of SQRD-1 and SQRD-2 and within a distance compatible with an electron transfer reaction during the simulation. Input files and evolution of protein RMSD, quinone-FAD distance and total energy during simulation time for each complex are available upon request.

**Figure 1:**
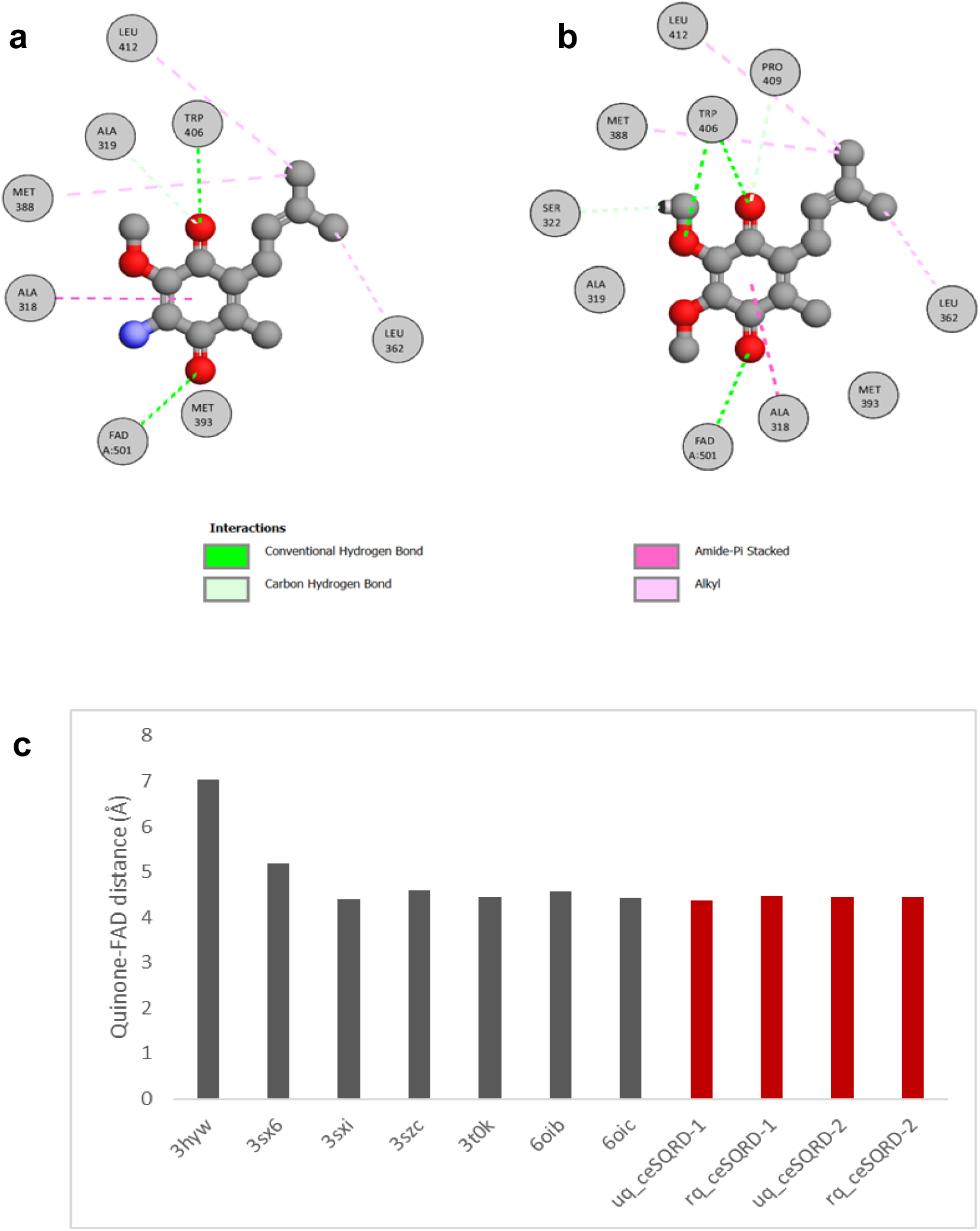
*C.elegans* SQRD-1 can accommodate both benzoquinones, UQ and RQ, at its active site. **a)** Interaction diagram of RQ docked in the homology model of SQRD-1. **b)** Interaction diagram of UQ docked in the homology model of SQRD-1. **c)** Comparison of the distance between the quinone ring and the imidic nitrogen of FAD in sulfide:quinone oxidoreductase structures deposited at PDB (grey) and the quinones ubiquinone (uq) and rhodoquinone (rq) docked in the homology models of *C.elegans* SQRD-1 and SQRD-2 (red). 3HYW: *Aquifex aeolicus* SQRD in complex with decylubiquinone; 3SX6: *Acidithiobacillus ferrooxidans* SQRD complexed with decylubiquinone, 3SXI: *Acidithiobacillus ferrooxidans* SQRD complexed with decylubiquinone, 3SZC: *Acidithiobacillus ferrooxidans* SQRD in complex with gold (I) cyanide, 3t0K: *Acidithiobacillus ferrooxidans* SQRD with bound trisulfide and decylubiquinone, 6OIB: human SQRD in complex with coenzyme Q, 6OIC: human SQRD in complex with coenzyme Q.

**Table 1.** Binding energy of RQ and UQ in complex with SQRD-1 and SQRD-2.

| **Benzoquinone** | **SQRD-1 binding energy (kcal/mol)** | **SQRD-2 binding energy (kcal/mol)** |
| --- | --- | --- |
| Ubiquinone | -7.4183 (GBVI) / -6.70<br>± 0.96 (KDeep) | -7,4253 (GBVI) / -6.63<br>± 0.53 |
| Rhodoquinone | -6.9301 (GBVI) / -6.76<br>± 0.87 (KDeep) | -7,1337 (GBVI) / -7.45<br>± 0.30 (KDeep) |

Interestingly, the gene models of *sqrd-1* and *sqrd-2* predict alternative transcripts and protein isoforms. Six isoforms are predicted for SQRD-1(a-f) and two for SQRD-2 (a and b) (**Figure S1**). All isoforms have the quinone binding site; however, the FAD-binding domain and the sulfide binding are absent in SQRD-1d and SQRD-1e isoforms. SQRD-1 b and SQRD-2 b isoforms possess N-terminal extensions, whose functional significance is yet to be characterized. In addition, three SQRD-1 isoforms (a, e and f) possess a 48-amino acid insertion that, according to modeling, would form an external loop not affecting the canonical folding of the protein.

We examined *sqrd-1(tm3378)* and *sqrd-2(ok3440)* mutant strains in the presence of sulfide in the liquid phase, using motility as a readout with *WMicrotracker* (**Figure 2**). Because the *WMicrotracker* is an open system, the sulfide concentration in solution decreases over time. We observed a decrease from 1.5 mM to 0 mM in 3 hours and from 0.5 mM to 0 in 2 hours (**Figure S3**). The *sqrd-1* mutant strain (*sqrd-1(tm3378)*) is more sensitive to sulfide than the wild-type N2 strain (**Figure 2**). The *sqrd-1(tm3378)* strain initially decreased its motility similarly to N2; however, once sulfur concentration decreased, the motility recovery of *sqrd-1(tm3378)* was slower than that of the N2 strain, suggesting that *sqrd-1* is part of an induced response to sulfide (**Figure 2**). In contrast to the *sqrd-1* mutant strain, the recovery of *sqrd-2* mutant strain (*sqrd-2(ok3440)*) was slightly earlier than that of the N2 strain (**Figure 2**). The results obtained suggest that, in addition to the previously described SQRD-1, SQRD-2 is also involved in the organism’s response to sulfide.

**Figure 2:**
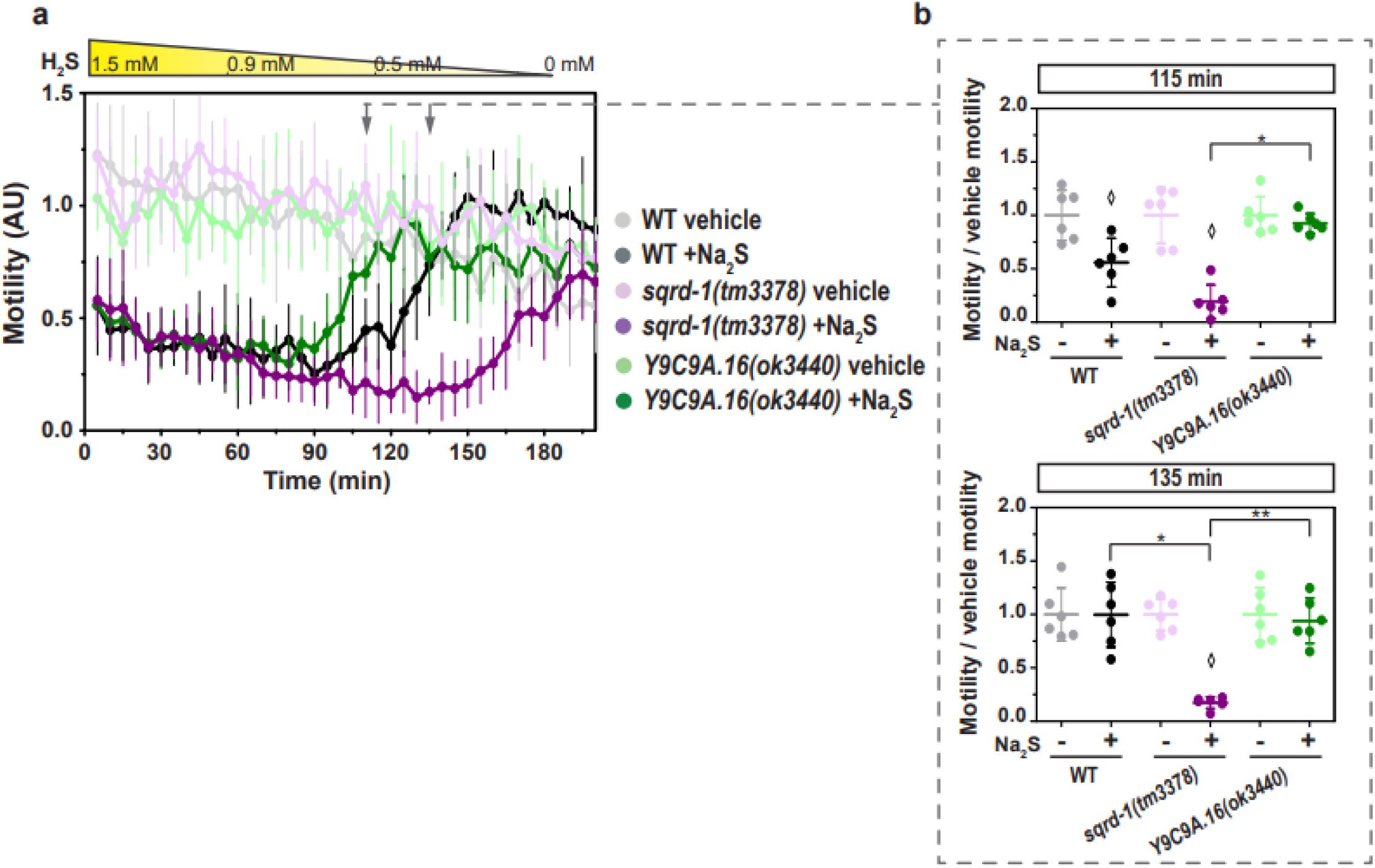
*C. elegans* possesses two sulfide quinone oxidoreductases. The motility parameter refers to the motility of a population of individuals in liquid media and was measured using the infrared tracking device *WMicrotracker*. Motility of *sqrd-1(tm3378)* and *Y9C9A.16(ok3440)* mutants and wild-type (WT) strains in the presence (+Na_2_S) or absence (vehicle) of sodium sulfide (Na_2_S) solution. Each point indicates the motility average of 6 wells (relative to the habituation, see methods) measured every 5 minutes for 200 minutes. The graph corresponds to a representative experiment with 6 wells per condition per strain (approximately 80 worms per well). Error bars indicate the standard deviation. At least three biological replicates were performed for each worm strain. The yellow triangle represents the decrease with time of the sulfide concentration in solution from 1.5 mM (initial concentration in the wells). Arrows represent the two time points shown in part b). AU: arbitrary units. **b**) For each strain (*sqrd-1(tm3378)*, *Y9C9A.16(ok3440)* and WT), the motility in the presence (+) or absence (-) of sulfide (Na_2_S) was normalized to the mean of the motility in the absence of Na_2_S. Each data point represents the normalized motility of each one of the 6 wells for each condition (with or without Na_2_S). Different graphs correspond to the time points of 115 and 135 minutes of incubation. Kruskal-Wallis test were performed (115 min p(value)=2.4E-5) was performed, followed by Dunńs post hoc test. ANOVA test (135 min p(value)=3.6E-9) was performed, followed by Tuckey pairwise. Asterisks indicate statistical differences between the strains indicated (115 min *p=2.5E-3 and 135 min *p=1.2E-6, **p=6.0E-6). Diamonds indicate statistical differences between sulfide and control vehicle (without sulfide) for each strain.

### RQ is essential for survival in sulfide

We next examined the relevance of RQ and UQ in H2S response. Since KYNU-1 is essential for RQ synthesis(44), we tested the *kynu-1(tm4924)* loss-of-function mutant for its response to H2S. This strain was highly sensitive to H2S, compared to the wild-type strain (**Figure 3a-b**). Importantly, almost all *kynu-1(tm4924)* animals died after 3 hours in H2S at 1.5 mM (94 % dead worms, **Figure S4**), whereas most wild-type worms were alive under these conditions. In contrast to *kynu-1(tm4924)*, the null-mutant strain *clk-1(qm30)*, unable to synthesize UQ, showed a complete motility recovery as the wild-type when H2S concentration decreased. Yet, the drop and recovery of motility in this mutant was more pronounced than that of the wild-type strain (**Figure 3a-b**). The same effect was observed with 0.5 mM of H2S (**Figure S4**). These results indicate that in these sulfide conditions, RQ was even more important than endogenous UQ for motility recovery and survival. Because the OP50 diet contains bacterial UQ, we then examined the sulfide response of the *clk-1 (qm30)* mutant strain fed during 20 hours with OP50 and afterwards with the UQ-less *E. coli* strain GD1 (feeding with GD1 from L1 result in larval arrest). The *clk-1* mutant strain fed with UQ-less diet showed a motility recovery similar to the results obtained using OP50 as food (**Figure 3c**). These results indicate that RQ is more relevant than UQ in the sulfide response, independently of the source of UQ.

**Figure 3:**
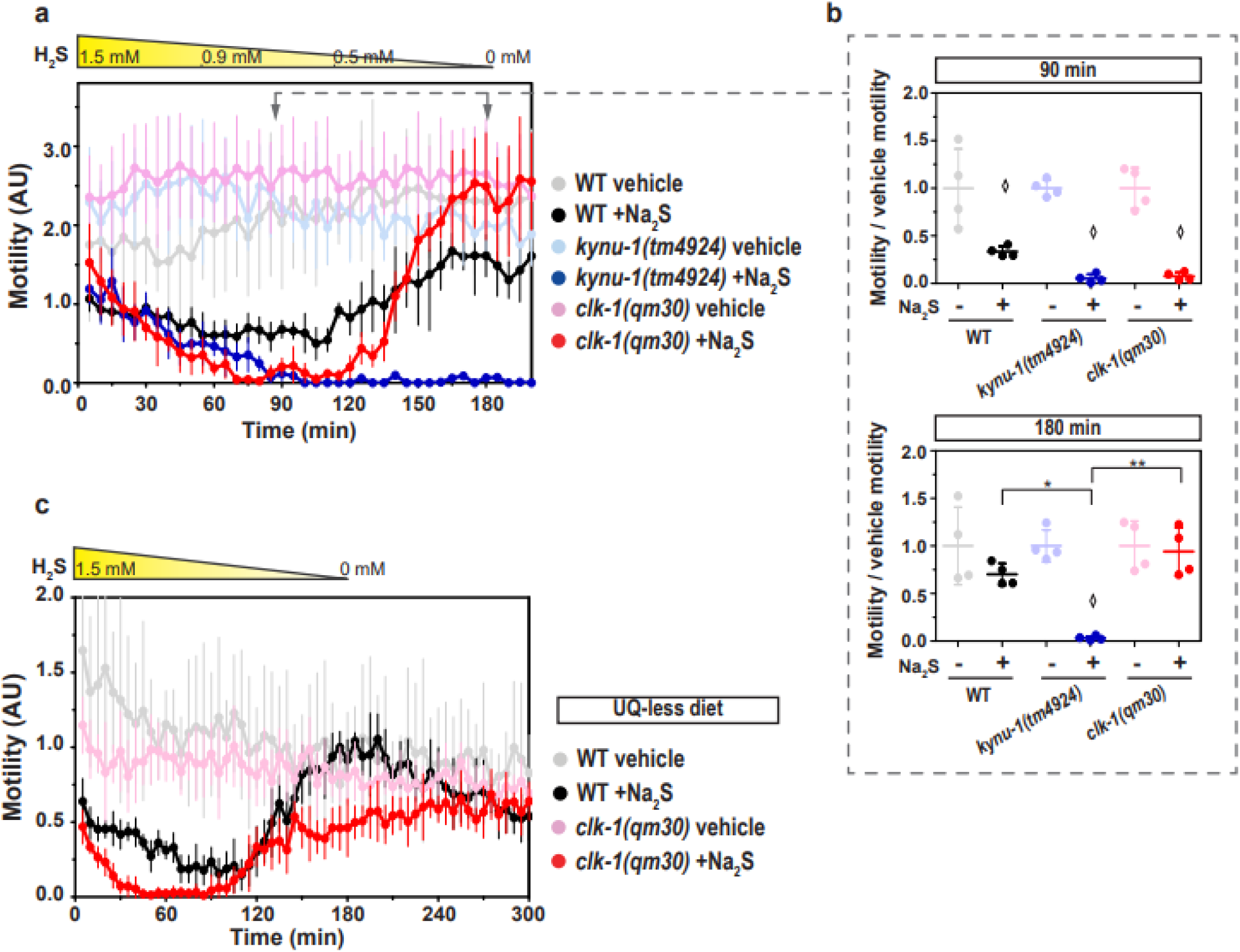
The rhodoquinone-deficient mutant strain (*kynu-1(tm4924)*) does not recover from sulfide challenge in contrast to the ubiquinone-deficient mutant strain (*clk-1(qm30)*). The motility parameter refers to the movement of a population of individuals in liquid media and was measured using the infrared tracking device *WMicrotracker*. **a**) Motility of *kynu-1(tm4924)* and *clk-1(qm30)* mutants and wild-type (WT) strains in the presence (+Na_2_S) or absence (vehicle) of a sodium sulfide (Na_2_S) solution. Each point indicates the motility average of 4 wells (relative to the habituation, see methods) measured every 5 minutes for 200 minutes. The graph corresponds to a representative experiment with 4 wells per condition per strain (approximately 80 worms per well). Error bars indicate the standard deviation. At least three biological replicates were performed for each worm strain. The yellow triangle represents the decrease of the sulfide concentration in solution from 1.5 mM. Arrows represent the two time points shown in part **b)**. AU: arbitrary units. **b)** For each strain (*kynu-1(tm4924)*, *clk-1(qm30)* and WT), the motility in the presence (+) or absence (-) of sulfide (Na_2_S) was normalized to the mean of the motility in the absence of Na_2_S. Each data point represents the normalized motility of each one of the 4 wells for the two conditions (with or without Na_2_S). Different graphs correspond to the time point of 90 and 180 minutes of incubation. Welch test (90 min p=1,83E-5 and 180min p=1.04E-4) were performed, followed by Tukeýs pairwise test. Asterisks indicate statistical differences (180min:*p=6.8E-3 and **p=1.5E-4). Diamonds indicate statistical differences for each strain between sulfide and their control without sulfide. **c)** Motility assay (with *WMicrotracker*) using worm strains (*clk-1(qm30)* and WT) that were fed with the UQ-deficient bacteria *Escherichia coli* GD1 strain. Motility of *clk-1(qm30)* mutant and WT strains in the presence (+Na_2_S) or absence (vehicle) of sodium sulfide (Na_2_S) solution. The points indicate the average of the motilities of 6 wells relative to the motility measured only with vehicle, every 5 minutes for 300 minutes. The graph corresponds to a representative experiment with 6 wells per condition per strain (approximately 80 worms per well). Error bars indicate the standard deviation. Three biological replicates were performed with similar results.

COQ-2 is essential for RQ and UQ biosynthesis(45). Organisms that synthesize RQ possess two isoforms of this enzyme (COQ-2A and COQ-2E), derived from mutually exclusive alternative splicing(45). COQ-2E, which is absent in organisms that do not synthesize RQ, has been shown to be essential for RQ biosynthesis. We examined the two COQ-2 isoforms mutants under sulfide conditions. Consistent with the results obtained with *kynu-1(tm4924)* and *clk-1(qm30)* mutant strains, the deletion mutant in exon 6e (*coq-2Δ6e*), was more sensitive to H2S than both the wild-type and the COQ-2a mutant strains (**Figure S5**).

We then analyzed the *kynu-1(tm4924)* mutant strain exposed to gaseous H2S (2 μM) for 20 hours and quantified the worm progeny. In the presence of H2S, *kynu-1(tm4924)* mutant animals showed a decreased offspring whereas the wild-type strain was unaffected (**Figure 4a-b**). Importantly, the expression of the *kynu-1* wild-type allele in the *kynu-1(tm4324)* mutant strain restored the progeny size in H2S, confirming the role of KYNU-1 in the H2S response.

**Figure 4:**
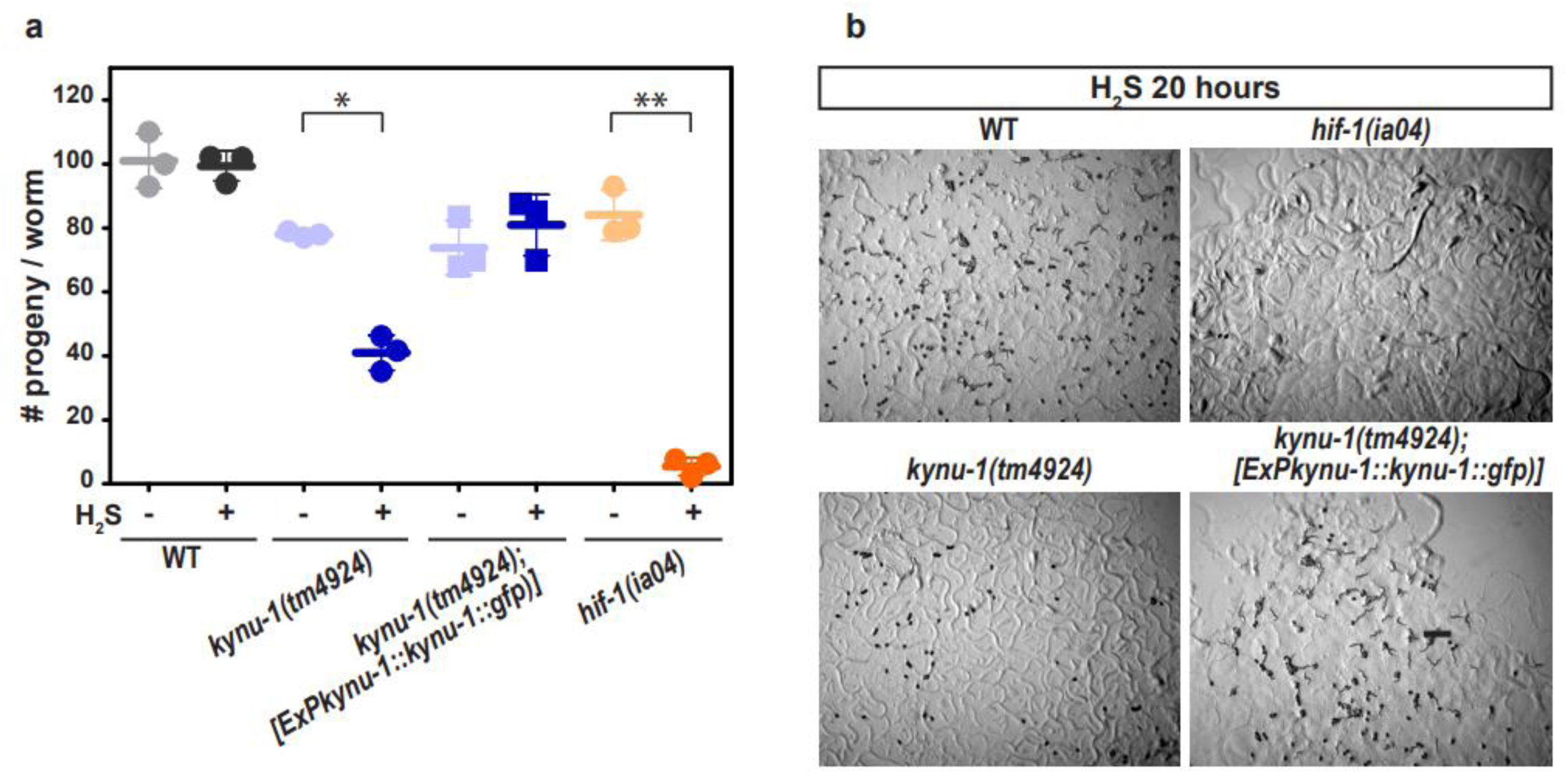
Sulfide affects the offspring size in the rhodoquinone-deficient mutant strain *kynu-1(tm4924)*. **a)** First day adults were exposed to a sulfide atmosphere (+H_2_S) or room air (-H_2_S) for 20 hours, and the number of progeny (embryos and L1) was scored. Each data point represents the mean of the number of offspring per adult worm obtained in an independent experiment (6 animals per condition per replicate were used). WT, wild-type N2 strain. Error bars represent the standard error of the mean. Means of three independent experiments are shown. ANOVA test was performed (p=8.5E-11) and subsequent Tukeýs pairwise test. Asterisks indicate statistical differences (*p=9E-5 and **p=5.6E-9). **b)** Representative images of the progeny of 6 adult worms after 20 hours in the H_2_S chamber of the following strains: wild-type N2, *kynu-1(tm4924), hif-1(ia04)* and *kynu-1(tm4924);ExPkynu-1::kynu-1::gfp*. Scale bar denote 500 µm. HIF-1 is required for the sulfide response and *hif-1(ia04)* mutant worms do not survive to 15 ppm of H_2_S (Budde, M., 2011).

Altogether, these results indicate that RQ is required for the organismal response to sulfur.

### RQ plays a role in the organismal response to pathogenic bacteria

We examined the RQ deficient mutant strains in response to *P. aeruginosa PAO1*, a bacterial strain that has been described to paralyze and kill the worm by cyanide production(10, 12). Cyanide detoxification leads to sulfide generation through CYSL-2(11). Furthermore, it has been shown that RQ confers an advantage for cyanide recovery(23). Therefore, RQ could confer an advantage to *C. elegans* response to *P. aeruginosa PAO1*. We found that the *kynu-1(tm4924)* strain was slightly more sensitive to the pathogenic bacteria than the wild-type strain (**Figure 5**). In contrast to *kynu-1(tm4924)*, the *clk-1(qm30)* mutant strain was similar to wild-type worms in response to the pathogenic bacteria PAO1 for three hours (**Figure 5**). Further exposure to this bacterial strain, led to motility decrease for all strains. Consistent with these results, the *coq-2Δ6e* was slightly more sensitive to the PAO1 bacteria than the wild-type strain (**Figure S6**). No difference was observed among the strains in the presence of the food bacterium *Escherichia coli* OP50 (**Figure S7**). These results highlight a putative role for RQ in the response to HCN-producing bacteria.

**Figure 5:**
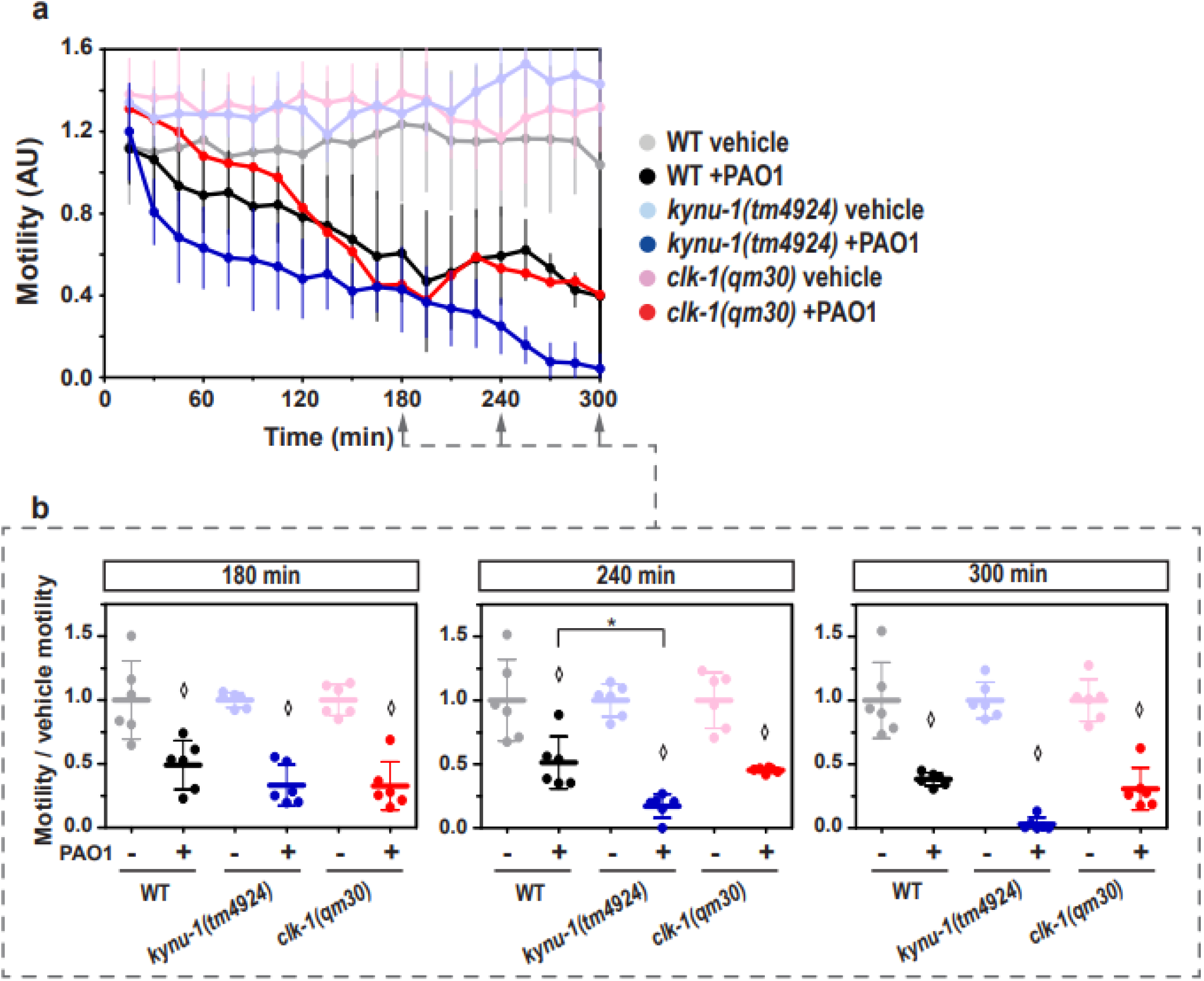
The rhodoquinone-deficient mutant strain *kynu-1(tm4924)* is more sensitive to *Pseudomonas aeruginosa* PAO1 strain than the ubiquinone-deficient mutant strain (*clk-1(qm30)*). The motility parameter refers to the movement of a population of individuals in liquid media and was measured using the infrared tracking device *WMicrotracker*. **a**) Motility of *kynu-1(tm4924)* and *clk-1(qm30)* mutants and wild-type (WT) strains in the presence (+PAO1) or absence (vehicle) of a liquid culture of *Pseudomonas aeruginosa* PAO1. Each point indicates the motility average of 4 wells (relative to the habituation, see methods) measured every 15 minutes for 300 minutes. The graph corresponds to a representative experiment with 6 wells per condition per strain (approximately 80 worms per well). Error bars indicate the standard deviation. At least three biological replicates were performed for each worm strain. **b**) For each strain (*kynu-1(tm4924)*, *clk-1(qm30)* and WT), the motility in the presence (+) or absence (-) of *P. aeruginosa* PAO1 was normalized to the mean of the motility in the absence of the bacteria. Each data point represents the normalized motility of each one of the 6 wells for the two conditions (with or without bacteria). Different graphs correspond to the time point of 180, 240 and 300 minutes of incubation. ANOVA test (180min p=9E-9) and Welch test (240 min p=7,4E-7) were performed, followed by Tukeýs pairwise test. Kruskal-Wallis test (300 min p=1,7E-5) were performed, followed by Dunńs post hoc test. The asterisk indicates statistical differences (*p=0,04). Diamonds indicate statistical differences for each strain between the two conditions (bacteria and vehicle).

### RQ does not play a role in oxidative stress

Ubiquinol, the reduced form of UQ, is a known lipid-soluble antioxidant for which there exist enzymic mechanisms that can regenerate its oxidized form, including SQRD/SQOR. In a halted ETC (e.g. cyanide or sulfide conditions) ubiquinol cannot be regenerated. Alternative pathways might, therefore, enhance an organism’s resistance to oxidative stress under such conditions, however it is unknown whether RQ can also function as an antioxidant, in a similar manner to UQ. In order to study a possible role of RQ in response to oxidative stress, the *kynu-1(tm4924)* mutant strain was exposed to paraquat, a chemical that generates reactive oxygen species mainly through a mechanism that involves complex I and cause cellular damage via lipid peroxidation(46). While the UQ-deficient mutant strain (*clk-1(qm30)*) was more sensitive to paraquat than the WT strain (**Figure 6**), the RQ-deficient mutant strain was not, and showed a similar profile to the WT strain in response to paraquat. Thus, in contrast to UQ, RQ has no evident role in the defense against oxidative stress.

**Figure 6:**
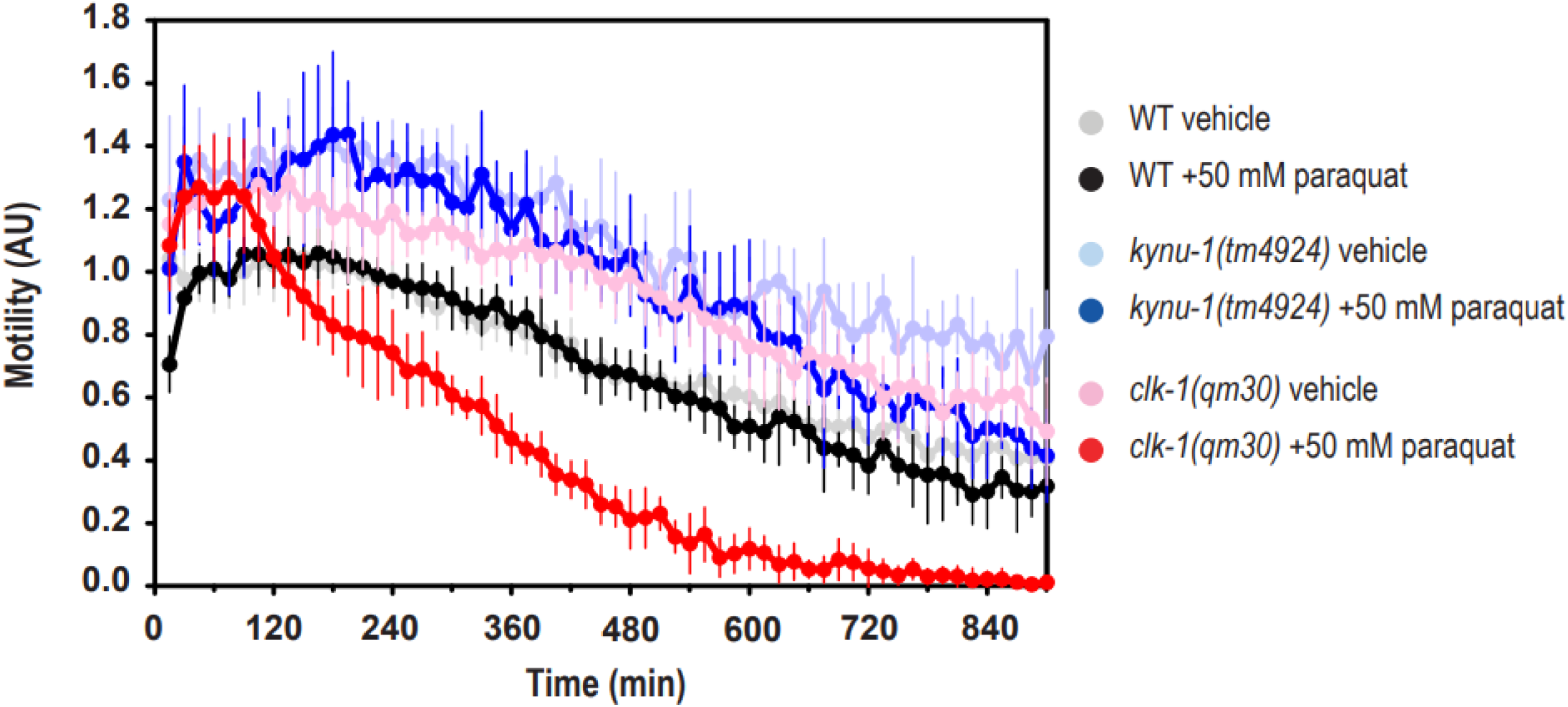
In contrast to the RQ-deficient mutant strain (*kynu-1(tm4924)*), the UQ-deficient mutant strain (*clk-1(qm30)*) was more sensitive to paraquat than the wild-type strain. The motility parameter refers to the movement of a population of individuals in liquid media and was measured using the infrared tracking device *WMicrotracker*. **a**) Motility of *kynu-1(tm4924)* and *clk-1(qm30)* mutants and wild-type (WT) strains in the presence (+paraquat) or absence (vehicle) 50 mM of paraquat. Each point indicates the motility average of 4 wells (relative to the habituation, see methods) measured every 15 minutes for 900 minutes. The graph corresponds to a representative experiment with 6 wells per condition per strain (approximately 80 worms per well). Error bars indicate the standard deviation. Three biological replicates with similar results were performed for each worm strain.

## Discussion

RQ is a lipidic electron carrier of an alternative mitochondrial ETC that constitutes a biochemical adaptation present in some animal lineages that face hypoxia (such as helminths, mollusks, and *C. elegans*)(19, 20). In this study, we demonstrate that the RQ-dependent ETC also serves a key role in a RQ-dependent sulfide oxidation pathway essential for sulfide detoxification.

In order to determine the role of RQ in sulfide *C. elegans* survival, we tested mutant strains unable to synthesize RQ (*kynu-1(tm4924)* and *coq-2(Δ6e)* strains(23, 44)) in the presence of sulfide. The two RQ-deficient mutant strains were highly sensitive to H2S. Indeed, RQ-less and UQ-plus worms fed with OP50, an UQ-containing diet, did not recover from sulfide (**Figure 3** and **Figure S5**). In contrast, UQ-less and RQ-plus worms (*clk-1(qm30)* and *coq-2(Δ6a)*) did recover from sulfide conditions when fed with either UQ-plus or UQ-less bacterial diet (**Figure 3** and **Figure S5**). This result indicates a more relevant role for RQ than UQ in worm survival under sulfide conditions. The essentiality of RQ can be attributed to a functional sulfide alternative oxidation pathway, operative when complex IV is inhibited by sulfide. We propose the model depicted in **Figure 7a**, contrasting the alternative sulfide oxidation ETC pathway with the canonical one. In this alternative ETC, SQRD oxidizes sulfide and reduces RQ, which in turn is regenerated by complex II which function as fumarate reductase, accepting electrons from reduced RQ. It is worth mentioning that it has been shown by Kumar, R. et al 2022(25) that mammalian complex II can function in reverse, oxidizing UQH2 to UQ allowing sulfide to be detoxified when complex IV is inhibited; yet, the standard redox potential of RQ would provide a much broader range of substrate concentrations to allow complex II to function as fumarate reductase.

**Figure 7:**
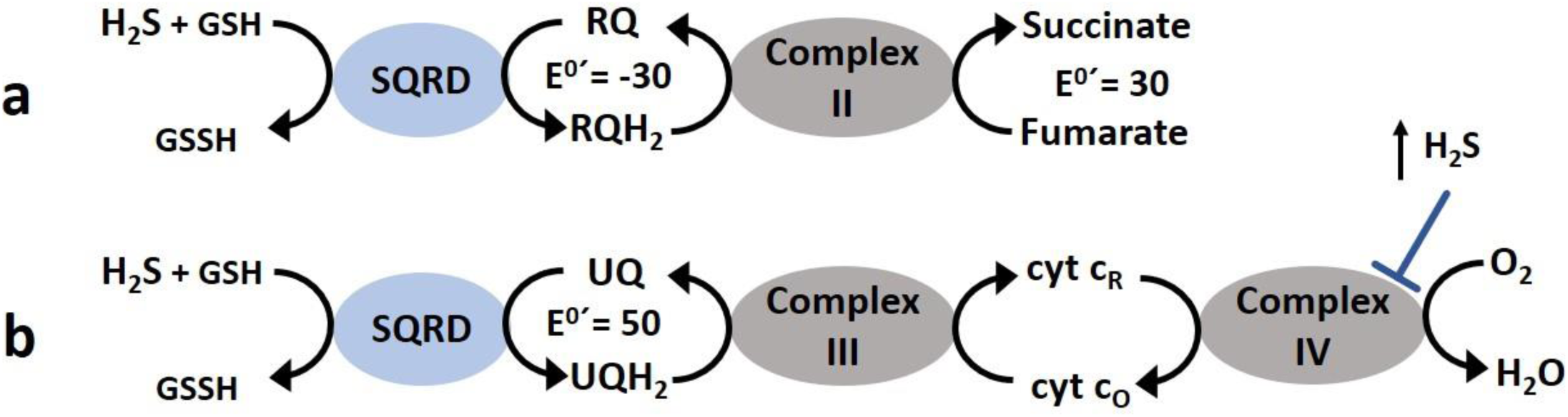
RQ- and UQ-dependent sulfide oxidation ETC. Based on our results, we propose the RQ-dependent model depicted **a)**, contrasting the canonical sulfide oxidation ETC pathway depicted in **b)**. In the alternative sulfide oxidation ETC, complex II functions as a fumarate reductase, allowing RQ to be recycled sustaining sulfide oxidation. Since complex IV is inhibited at high sulfide concentrations, this RQ-dependent pathway confers an advantage over the canonical pathway at toxic sulfide concentrations. Although it has been described that complex II can function as fumarate reductase with UQ, the standard reduction potential of RQH_2_/RQ, lower than that of fumarate/succinate allows sulfide oxidation over a broader range of concentrations. The standard reduction E^0^′ potential of rhodoquinone/rhodoquinol, ubiquinone/ubiquinol and fumarate/succinate are expressed in millivolts.

Remarkably, mutants unable to synthesize RQ exhibited even greater sensitivity to sulfide compared to the *sqrd-1* mutant (Figure 2 and Figure 3). Because RQ can also accept electrons from complex I and transfer them to fumarate via complex II (**Figure 7b**) it would also help to maintain ATP/ADP, NADH/NAD^+^ and mitochondrial membrane potential homeostasis and to decrease the oxidative stress caused by a sulfide halted canonical ETC. Additional mechanisms of RQ contribution to sulfide detoxification and survival cannot be ruled out. Although RQ may prevent oxidative damage from occurring in the first place, decreasing the superoxide radical production by complex I, our results indicate that RQ cannot function as a direct antioxidant to ameliorate paraquat-induced oxidative stress (Figure 6).

Another toxin commonly encountered by *C. elegans* in its natural environment is hydrogen cyanide produced by pathogenic bacterial strains that paralyze and kill the worm(10, 12). Similar to sulfur, cyanide also inhibits complex IV. Furthermore, RQ is essential for worm survival in cyanide(23). The mechanism proposed for HCN detoxification in the worm involves the enzyme CYSL-2 that consumes cyanide and generates H2S. The H2S produced triggers a recovery response to both chemicals and is detoxified mainly by the sulfide oxidation pathway(11). Importantly, the RQ-less mutant strain was slightly more sensitive to the cyanide-producing bacteria *P. aeruginosa* PAO1 than both the wild type and the UQ-less mutant strains (**Figure 4 and Figure S6**).

Interestingly, in contrast to mammals, *C. elegans* possess two SQRD/SQOR genes, *sqrd-1* and *sqrd-2*, the latter initially annotated as a pseudogen (quoted in (11)). However, we showed that *sqrd-2* is expressed in *C. elegans*. In concordance with our result, transcriptionally active regions of the *sqrd-2* gene were described in a tiling-array project run as part of the modENCODE initiative(47). Unlike the *sqrd-1* gene, *sqrd-2* does not change its expression under H2S conditions according to microarrays studies(16). On the other hand, RNAseq data obtained using strains that overexpress HIF-1 showed a 1.3-1.9-fold change in *sqrd-2* expression, suggesting a regulation of *sqrd-2* expression in a HIF-1 dependent manner(48). Both SQRD-1 and SQRD-2 would be able to recognize RQ and UQ in their active sites; yet, the mutant strains exhibited different phenotypes in sulfide response. The *sqrd-1* mutant strain was more sensitive to sulfide than the wild-type N2 strain (**Figure 2**), confirming its previously described role in sulfide detoxification. In contrast, the *sqrd-2* mutant strain recovered earlier than the wild-type strain, suggesting that SQRD-2 would function as a negative regulator of the sulfide response. It is interesting to note that *sqrd* gene duplication occurs in other animals that face sulfide and/or hypoxia (**Figure S1**).

To sum up, we describe a new RQ function, key in the adaptive biochemical response to sulfide in *C. elegans*. Importantly, the alternative sulfide oxidation pathway described would confer an advantage in the presence of H2S and HCN, and hence to pathogenic bacteria producing these toxins. More broadly, the animal lineages well-adapted to sulfide-rich environments (nematodes, annelids and bivalves) synthesize RQ. This reinforces the concept that RQ is a key biochemical adaptation not only for hypoxia, but also for sulfide-rich environments.

### Experimental procedures

#### Strains and culture conditions

General methods for *C. elegans* culture and maintenance were performed according to reference(26). All chemical reagents, unless otherwise specified, were purchased from Sigma-Aldrich-Merck (St. Louis, MO). **Table 2** lists the *C. elegans* strains used in this study, detailing the genotype and source. The *Escherichia coli* OP50 strain used as *C. elegans* food was obtained from the Caenorhabditis Genetic Center (CGC). The *Pseudomonas aeruginosa* PAO1 was kindly provided by Dr. Colin Manoil (Universidad de Washington, USA) and the ubiquinone-deficient *E. coli* strain GD1 was kindly provided by Dr. Catherine Clarke (UCLA, USA).

**Table 2.** *C. elegans* strains used in this study, detailing the genotype and the source. CGC: *Caenorhabditis* Genetic Center and NBPJ: *C. elegans* National Bioresource Project of Japan.

| Strain | Genotype | Source |
| --- | --- | --- |
| Bristol N2 | Wild-type | CCG |
| TM3378 | *sqrd-1(tm3378) IV* | NBPJ |
| RB2485 | *Y9C9A.16(ok3440) IV* | CGC |
| TM4924 | *kynu-1(tm4924) X* | NBPJ |
| MQ130 | *clk-1(qm30) III* | CGC |
| IH25 | *kynu-1(tm4924)X*;<br>*Ex[Pkynu-1::kynu1::gfp,*<br>*pRF4]* | Institut Pasteur de Montevideo.<br>Roberts Bucetta, P. *et al.*, 2019 |
| PHX1715 | *coq-2 (syb1715) III* | Suny Biotech Co., Ltd (Fuzhou City, China). Tan, J. H. *et al.*, 2020 |
| PHX1721 | *coq-2 (syb1721) III* | Suny Biotech Co., Ltd (Fuzhou City, China). Tan, J. H. *et al.*, 2020 |
| ZG31 | *hif-1(ia4) V* | CGC |

#### Sulfide solutions and quantification of the concentration

Crystals of Na2S·9H2O were stored in a desiccator until use. Solutions containing sulfide were prepared in ultrapure water sealed vials. The sulfide concentration in solution was determined by the methylene blue colorimetric assay described in Siegel, L. M, 1964(27) using an absorbance microplate reader Varioskan^TM^. The light path was determined as 0.58 cm for a 96 well plate and a total solution volume of 200 µL.

The gaseous H2S for the sulfide chamber was generated using a solution of Na2S·9H2O in chloride acid (1.31E-5 moles of Na2S in 8 mL of HCl 50 mM) in a bohemian glass inside a hermetic chamber (1.44 L volume) and gentle stirring (40 rpm). The gaseous H2S concentration was measured by the methylene blue method, mentioned above, extracting 2 mL of gas with a gas-tight Hamilton syringe and bubbling the gas in the alkaline zinc acetate solution of the assay.

#### Toxicity tests using the tracking device WMicrotracker ONE

The toxicity test in liquid media was determined using the tracking device *WMicrotracker^TM^ ONE* (PhylumTech, Argentina). The method used to determine worm motility is described in detail in(28). Briefly, the system detects motility through the interference of an array of infrared light beams, caused by worm movement. The readout is counts per unit of time (5 or 15 min in our experiments). Each count represents the interruption of the infrared light beam by worms. Experiments were performed in 96 well plates, using 80-100 synchronized L4 animals per well in a final volume of 100 µL. Six wells per condition per strain were used in each experiment (technical replicates). Experiments were repeated at least three times (biological replicates). In all cases, the counts per well at different times are normalized by the counts before adding the compound of interest or M9 buffer vehicle (basal counts or “habituation”). This normalization corrects the small differences that may exist in the number of worms per well. All the assays included the wild-type strain and vehicle for each strain as controls.

For the experiments performed with GD1 as worm food, synchronized L1 larvae were placed in OP50 seeded NGM plates for 24 hours and then washed three times in M9 and transferred to GD1 seeded NGM plates. The initial growth with OP50 is needed because *clk-1* worms fed on GD1 from L1 showed larval arrest. Because the different developmental time between the wild-type N2 and the *clk-1(qm30)* strains, the wild-type was incubated at 15°C and the *clk-1* mutant at 20°C until the motility assay. The *WMicrotracker* assays were performed using L4 worms.

#### Survival and progeny in sulfide atmosphere

For survival assays and progeny quantification, 6-day adults *C. elegans* were placed on a 3-cm NGM plate seeded with OP50 bacteria. The plates were exposed to a H2S in room air (RA) atmosphere for 20 hours and survival was assessed immediately after removal from the H2S/RA atmosphere. Animals were considered dead if they did not respond to tapping the plate or the worm touch with the wire pick. Additionally, after H2S exposure, the number of eggs and larvae was scored in each plate. The maximal H2S concentration obtained within the chamber was 2 µM. For each strain, identical plates without sulfide (only RA) were included as controls. Experiments were conducted at 20°C.

#### Statistical analysis

Normality and variance homogeneity were determined by Shapiro-Wilk and Levenés tests, respectively, with a 5% of significance level. Normal data were analyzed by ANOVA test and subsequent Tukey’s test for pairwise comparisons. Samples with unequal variances were compared using Welch F test and Tukey’s test for pairwise comparisons. Non-parametric data were analyzed using Kruskal-Wallis test and Dunńs post-hoc test for pairwise comparisons.

#### RNA isolation and Y9C9A.16 PCR amplification

RNA was extracted from ∼50-mg N2 worm pellets. For that 4000 synchronized L1s were grown on NGM plates at 20 °C to adulthood. For each experiment, ∼10000 adult worms were harvested and washed three times with 18 megohm water to obtain pellets for extraction (∼50 mg) and frozen at -80°C. The thawed pellet was ground in a Bullet Blender (Next Advance, USA) with 50 mg of zirconium oxide beads of 50 mm (Next Advance) for 2 min at power 8. Ground worms were used for RNA isolation using the QiAmp Viral RNA mini kit (Qiagen) according to the manufacturer’s instructions. The RNA concentration was quantified using a NanoDrop spectrophotometer (Thermo Fisher Scientific) and the RNA was treated with a DNase I enzyme (Roche).

For the cDNA synthesis oligodT was used and the retrotranscription was performed using the Superscript IV reverse transcriptase (Thermo Fisher Scientific) enzyme. The PCR amplification of Y9C9A.16 gene fragment was performed using the specific primers gtcgacagttttcaagtgcg and ctcgaaaacttccaagtggc.

#### Homology modeling

SQRD-1 and SQRD-2 sequences were retrieved from Wormbase and a pdbBLAST search was performed in USCF ChimeraX(29). Human sulfide:quinone oxidoreductase (SQOR; pdb code 6OIB) was identified as the most similar sequence with its experimental structure determined (similarity 59%) and used as a template for homology modeling. Homology models were generated using Phyre2(30), Swissmodel, Modeller(31), iTasser(32) and MOE(33). An energy minimization was performed on all models using Amber14:EHT(34) forcefield implemented in MOE. RMSD between the model and the template, QMean score(35) and visual inspection were used to assess the quality of the models generated. The MOE model was selected for docking experiments given its good QMean score and the better loop modeling.

#### Docking

We previously optimized a docking protocol in MOE for the Q site of *C. elegans* mitochondrial_complex II(36). The same protocol was validated using the human SQOR (pdb code 6OIB) using the co-crystallized ligand (ubiquinone). The RMSD value between the docked and the co-crystallized ligand was 0.137 Å. The protocol was then applied for ubiquinone and rhodoquinone docking with *C. elegans* SQRD-1 and SQRD-2 models. GBVI and KDeep score functions were employed as binding energy estimators.

#### Molecular dynamics

The four complexes obtained from docking experiments (SQRD-1b:UQ, SQRD-1b:RQ, SQRD-2b:UQ and SQRD2b:RQ) and the template crystal 6OIC were prepared using the QwikMD module(37, 38) of VMD 1.9.4 with the default settings except for the duration of each production stage, which was increased to 5000000 steps (10 ns). Five production stage were concatenated to give a total simulation time of 50 ns for each complex and 6OIC. Atom typing and CHARMM(39) forcefield parameter assignation for hydrogen sulfide, FAD and quinones were performed using CGenFF(40) server. Calculations were run in the NAMD 3.0(41) alpha version using CUDA acceleration in a PC with a 12th Gen Intel® Core™ i5-12400 × 12 processor and an NVIDIA GeForce RTX 4080/PCIe/SSE2 GPU with Ubuntu 22.04 LTS as the operating system. VMD was used for structural analysis and visualization.

#### Paraquat assay

A 0.1 M methyl viologen dichloride hydrate solution was prepared in ultrapure water and was added to an equal volume of M9 buffer containing worms in a 96-well plate. Worm motility assays were performed using the *WMicrotracker*.

#### Pseudomonas aeruginosa PAO1 assays

*P. aeruginosa* were growth in King B medium described by Bio-Rad, supplemented with glycine (4.4 g/L) and FeCl3 (100 µM) to promote cyanide production as described in(42). An overnight culture was diluted in the same medium to obtain the same OD (at 600 nm) than the *E. coli* OP50 culture used as control.

## Supporting information

This article contains supporting information

## Acknowledgements

We would like to thank Dr. Ernesto Cuevasanta (Laboratorio de Fisicoquímica Biológica, Facultad de Ciencias, Universidad de la República, Uruguay) for technical assistance in sulfide determination; Caenorhabditis Genetic Center (CGC) for *C. elegans* mutant strains; Dr. Colin Manoil (Universidad de Washington, USA) and Dr. Catherine Clarke (UCLA, USA) for *P. aeruginosa* and *E. coli* strains.

## References

1. Nicholls, P., and Kim, J. K. (1982) Sulphide as an inhibitor and electron donor for the cytochrome c oxidase system. Can. J. Biochem. 60, 613– 623

2. Filipovic, M. R., Zivanovic, J., Alvarez, B., and Banerjee, R. (2018) Chemical Biology of H2S Signaling through Persulfidation. Chem. Rev. 118, 1253–1337

3. Bailly, X., and Vinogradov, S. (2005) The sulfide binding function of annelid hemoglobins: Relic of an old biosystem? J. Inorg. Biochem. 99, 142–150

4. Broman, E., Bonaglia, S., Holovachov, O., Marzocchi, U., Hall, P. O. J., and Nascimento, F. J. A. (2020) Uncovering diversity and metabolic spectrum of animals in dead zone sediments. *Commun*. Biol. 3, 1–12

5. Sun, Y., Wang, M., Zhong, Z., Chen, H., Wang, H., Zhou, L., Cao, L., Fu, L., Zhang, H., Lian, C., Sun, S., and Li, C. (2022) Adaption to hydrogen sulfide-rich environments: Strategies for active detoxification in deep-sea symbiotic mussels, Gigantidas platifrons. Sci. Total Environ. 804, 150054

6. Salinas, G., Langelaan, D. N., and Shepherd, J. N. (2020) Rhodoquinone in bacteria and animals: Two distinct pathways for biosynthesis of this key electron transporter used in anaerobic bioenergetics. Biochim. Biophys. Acta - Bioenerg. 10.1016/j.bbabio.2020.148278

7. Libiad, M., Yadav, P. K., Vitvitsky, V., Martinov, M., and Banerjee, R. (2014) Organization of the human mitochondrial hydrogen sulfide oxidation pathway. J. Biol. Chem. 289, 30901–30910

8. Jackson, M. R., Melideo, S. L., and Jorns, M. S. (2012) Human sulfide:Quinone oxidoreductase catalyzes the first step in hydrogen sulfide metabolism and produces a sulfane sulfur metabolite. Biochemistry. 51, 6804–6815

9. Theissen, U., Hoffmeister, M., Grieshaber, M., and Martin, W. (2003) Single eubacterial origin of eukaryotic sulfide:quinone oxidoreductase, a mitochondrial enzyme conserved from the early evolution of eukaryotes during anoxic and sulfidic times. Mol. Biol. Evol. 20, 1564– 1574

10. Gallagher, L. A., and Manoil, C. (2001) Pseudomonas aeruginosa PAO1 Kills Caenorhabditis elegans by Cyanide Poisoning. J. Bact. 183, 6207–6214

11. Budde, M. W., and Roth, M. B. (2011) The response of caenorhabditis elegans to Hydrogen Sulfide and Hydrogen Cyanide. Genetics. 189, 521–532

12. Darby, C., Cosma, C. L., Thomas, J. H., and Manoil, C. (1999) Lethal paralysis of Caenorhabditis elegans by Pseudomonas aeruginosa. Proc. Natl. Acad. Sci. U. S. A. 96, 15202–15207

13. Wang, B., Pandey, T., Long, Y., Delgado-Rodriguez, S. E., Daugherty, M. D., and Ma, D. K. (2022) Co-opted genes of algal origin protect C. elegans against cyanogenic toxins. Curr. Biol. 32, 4941–4948.e3

14. Ma, D. K., Vozdek, R., Bhatla, N., and Horvitz, H. R. (2012) CYSL-1 Interacts with the O 2-Sensing Hydroxylase EGL-9 to Promote H 2S-Modulated Hypoxia-Induced Behavioral Plasticity in C. elegans. Neuron. 73, 925–940

15. Vozdek, R., Hnízda, A., Krijt, J., Šerá, L., and Kožich, V. (2013) Biochemical properties of nematode O-acetylserine(thiol)lyase paralogs imply their distinct roles in hydrogen sulfide homeostasis. Biochim. Biophys. Acta - Proteins Proteomics. 1834, 2691–2701

16. Miller, D. L., Budde, M. W., and Roth, M. B. (2011) HIF-1 and SKN-1 coordinate the transcriptional response to hydrogen sulfide in Caenorhabditis elegans. PLoS One. 10.1371/journal.pone.0025476

17. Budde, M. W., and Roth, M. B. (2010) Hydrogen Sulfide Increases Hypoxia-Inducible Factor-1 Activity Independently of von Hippel-Lindau Tumor Suppressor-1 in C. elegans. Mol. Biol. Cell. 21, 212–217

18. Horsman, J. W., Heinis, F. I., and Miller, D. L. (2019) A novel mechanism to prevent H2S toxicity in caenorhabditis elegans. Genetics. 213, 481–490

19. Sato, M., and Ozawa, H. (1969) Occurrence of Ubiquinone and Rhodoquinone in Parasitic Nematodes, Metastrongylus elongatus and Ascaris lumbricoides va.suis. J. Biochem. 65, 861–867

20. Allen, P. C. (1973) Helminths: Comparison of their rhodoquinone. Exp. Parasitol. 34, 211–219

21. Tielens, A. G. M., and Van Hellemond, J. J. (1998) The electron transport chain in anaerobically functioning eukaryotes. Biochim. Biophys. Acta - Bioenerg. 1365, 71–78

22. Van Hellemond, J. J., Klockiewicz, M., Gaasenbeek, C. P. H., Roos, M. H., and Tielensi, A. G. M. (1995) Rhodoquinone and complex II of the electron transport chain in anaerobically functioning eukaryotes. J. Biol. Chem. 270, 31065–31070

23. Del Borrello, S., Lautens, M., Dolan, K., Tan, J. H., Davie, T., Schertzberg, M., Spensley, M. A., Caudy, A. A., and Fraser, A. G. (2019) Rhodoquinone biosynthesis in 1 C.elegans requires precursors generated by the kynurenine pathway. Elife. 8, 1–21

24. Erabi, T., Higuti, T., Tomisaburo Kakuno, -l, Yamashita, J., Tanaka, M., and Horio, T. (1975) Polarographic Studies on Ubiquinone-10 and Rhodoquinone Bound with Chromatophores from Rhodospirillum rubrum

25. Kumar, R., Landry, A. P., Guha, A., Vitvitsky, V., Lee, H. J., Seike, K., Reddy, P., Lyssiotis, C. A., and Banerjee, R. (2022) A redox cycle with complex II prioritizes sulfide quinone oxidoreductase dependent H2S oxidation. J. Biol. Chem. 10.1016/j.jbc.2021.101435

26. Sulston, J. E., and Hodgkin, J. (1997) Methods. in The Nematode Caenorhabditis elegans (Cold Spring Harbor Monograph Series 17) (Wood, W. B. ed), pp. 587–606, Cold Spring Harbor, Boulder

27. Siegel, L. M. (1965) A direct microdetermination for sulfide. Anal. Biochem. 11, 126–132

28. Simonetta, S. H., and Golombek, D. A. (2007) An automated tracking system for Caenorhabditis elegans locomotor behavior and circadian studies application. J. Neurosci. Methods. 161, 273–280

29. Pettersen, E. F., Goddard, T. D., Huang, C. C., Meng, E. C., Couch, G. S., Croll, T. I., Morris, J. H., and Ferrin, T. E. (2021) UCSF ChimeraX: Structure visualization for researchers, educators, and developers. Protein Sci. 30, 70–82

30. Kelley, L. a, Mezulis, S., Yates, C. M., Wass, M. N., and Sternberg, M. J. E. (2015) Europe PMC Funders Group The Phyre2 web portal for protein modelling , prediction and analysis. Nat. Protoc. 10, 845–858

31. Kiefer, F., Arnold, K., Künzli, M., Bordoli, L., and Schwede, T. (2009) The SWISS-MODEL Repository and associated resources. Nucleic Acids Res. 37, 387–392

32. Xiaogen, Z., Zheng, W., Li, Y., Pearce, R., Zhang, C., Bell, E. W., Zhang, G., and Zhang, Y. (2022) I-TASSER-MTD: a deep-learning-based platform for multi-domain protein structure and function prediction. Nat. Protoc*. Vol.* 17, 2326–2353

33. 2022.02 Chemical Computing Group ULC Molecular Operating Environment (MOE)

34. D.A. Case J.T. Berryman, R.M. Betz, Q. Cai, D.S. Cerutti, T.E. Cheatham, III, T.A. Darden, R.E. Duke, H. Gohlke, A.W. Goetz, S. Gusarov, N. Homeyer, P. Janowski, J. Kaus, I. Kolossváry, A. Kovalenko, T.S. Lee, S. LeGrand, T. Luchko, R. Luo, B. Madej, K.M, V. B. (2014) The Amber Molecular Dynamics Package. Amber

35. Benkert, P., Tosatto, S. C. E., and Schomburg, D. (2008) QMEAN: A comprehensive scoring function for model quality assessment. Proteins Struct. Funct. Genet. 71, 261–277

36. Vairoletti, F., Paulino, M., Mahler, G., Salinas, G., and Saiz, C. (2022) Structure-Based Bioisosterism Design, Synthesis, Biological Evaluation and In Silico Studies of Benzamide Analogs as Potential Anthelmintics. Molecules. 10.3390/molecules27092659

37. Ribeiro, J. V., Bernardi, R. C., Rudack, T., Stone, J. E., Phillips, J. C., Freddolino, P. L., and Schulten, K. (2016) QwikMD - Integrative Molecular Dynamics Toolkit for Novices and Experts. Sci. Rep. 6, 1–14

38. Humphrey, W., Dalke, A., and Schulten, K. (1996) VMD: Visual Molecular Dynamics. J. Mol. Graph. 14, 33–38

39. K. Vanommeslaeghe, E. Hatcher, C. Acharya, S. Kundu, S. Zhong, J. Shim, E. Darian, O. Guvench, P. Lopes, I. Vorobyov, A. D. M. J. (2010) J Comput Chem - 2009 - Vanommeslaeghe - CHARMM general force field A force field for drug-like molecules compatible with.pdf

40. Vanommeslaeghe, K., and Mackerell, A. D. J. (2012) Automation of the CHARMM General Force Field (CGenFF) I: Bond Perception and Atom Typing. J. Chem. Inf. Model. 52, 3144–3154

41. Phillips, J. C., Hardy, D. J., Maia, J. D. C., Stone, J. E., Ribeiro, J. V., Bernardi, R. C., Buch, R., Fiorin, G., Hénin, J., Jiang, W., McGreevy, R., Melo, M. C. R., Radak, B. K., Skeel, R. D., Singharoy, A., Wang, Y., Roux, B., Aksimentiev, A., Luthey-Schulten, Z., Kalé, L. V., Schulten, K., Chipot, C., and Tajkhorshid, E. (2020) Scalable molecular dynamics on CPU and GPU architectures with NAMD. J. Chem. Phys. 10.1063/5.0014475

42. Bakker, A. W., and Schippers, B. (1987) Microbial cyanide production in the rhizosphere in relation to potato yield reduction and Pseudomonas SPP-mediated plant growth-stimulation. Soil Biol. Biochem. 19, 451–457

43. Mishanina, T. V., Yadav, P. K., Ballou, D. P., and Banerjee, R. (2015) Transient kinetic analysis of hydrogen sulfide oxidation catalyzed by human sulfide quinone oxidoreductase. J. Biol. Chem. 290, 25072– 25080

44. Roberts Buceta, P. M., Romanelli-Cedrez, L., Babcock, S. J., Xun, H., VonPaige, M. L., Higley, T. W., Schlatter, T. D., Davis, D. C., Drexelius, J. A., Culver, J. C., Carrera, I., Shepherd, J. N., and Salinas, G. (2019) The kynurenine pathway is essential for rhodoquinone biosynthesis in Caenorhabditis elegans. J. Biol. Chem. 294, 11047–11053

45. Tan, J. H., Lautens, M., Romanelli-Cedrez, L., Wang, J., Schertzberg, M. R., Reinl, S. R., Davis, R. E., Shepherd, J. N., Fraser, A. G., and Salinas, G. (2020) Alternative splicing of COQ-2 determines the choice between ubiquinone and rhodoquinone biosynthesis in helminths. bioRxiv

46. Cochemé, H. M., and Murphy, M. P. (2008) Complex I is the major site of mitochondrial superoxide production by paraquat. J. Biol. Chem. 283, 1786–1798

47. Mark B. Gerstein, Zhi John Lu, Eric L. Van Nostrand, Chao Cheng, Bradley I. Arshinoff, Tao Liu, Kevin Y. Yip, Rebecca Robilotto, Andreas Rechtsteiner, Kohta Ikegami, Pedro Alves, Aurelien Chateigner, Marc Perry, Mitzi Morris, Raymond K. Auerbach, Xin Feng, J. D. L. and R. H. W. (2011) Integrative Analysis of the Caenorhabditis elegans Genome by the modENCODE Project. Science (80-. ). 330, 1775–1787

48. Vora, M., Pyonteck, S. M., Popovitchenko, T., Matlack, T. L., Prashar, A., Kane, N. S., Favate, J., Shah, P., and Rongo, C. (2022) The hypoxia response pathway promotes PEP carboxykinase and gluconeogenesis in C. elegans. Nat. Commun. 10.1038/s41467-022-33849-x

