## Supporting information for "Rhodoquinone-dependent electron transport chain is essential for *C. elegans* survival in hydrogen sulfide environments"

Material included:

**Figure S1:** Alignment of sulfide:quinone oxidoreductases from *C. elegans* and other animal lineages that possess two SQRD genes with the human protein.

**Figure S2:** *C. elegans Y9C9A.16* (*sqrd-2*) gene model and mRNA PCR amplification product.

**Figure S3:** $H_2S$ concentration in solution decay over the time. $H_2S$ concentration in solution was determined using the colorimetric assay.

**Figure S4:** The rhodoquinone-deficient mutant strain (*kynu-1(tm4924)*) does not recover from sulfide challenge in contrast to the ubiquinone-deficient mutant strain (*clk-1(qm30)*).

**Figure S5:** The rhodoquinone-deficient mutant strain (*coq-2(syb1721)*, (*coq-2Δ6e*)) does not recover from sulfide challenge in contrast to the ubiquinone-deficient mutant strain (*coq-2(syb1715)*, (*coq-2Δ6a*)).

**Figure S6:** The rhodoquinone-deficient mutant strain *coq-2(syb1721)* (*coq-2Δ6e*) is more sensitive to *Pseudomonas aeruginosa* PAO1 strain than the wild-type strain.

**Figure S7:** Liquid culture of the food bacterium *Escherichia coli* OP50 does not affect differentially the motility of the worm mutant strains *kynu-1(tm4924)*, *clk-1(qm30)*, *coq-2(syb1721)(coq-2Δ 6e)*, in comparison with the wild-type strain.

```
SQRD-1_isoform_b                 --------------------------------MRTSTVVYGKHFKLLVVG
SQRD-1_isoform_c                 --------------------------------------------------
SQRD-1_isoform_a                 --------------------------------MRTSTVVYGKHFKLLVVG
SQRD-1_isoform_f                 --------------------------------------------------
SQRD-1_isoform_d                 --------------------------------------------------
SQRD-1_isoform_e                 --------------------------------------------------
Y9C9A.16_isoform_a               --------------------------------------------------
Y9C9A.16_isoform_b               --------------------------------MITSAVLHAKHFKLLVVG
Ascaris_suum_tr|F1L539           --------------------------------------------------
Ascaris_suum_tr|F1L970           -------------------------------MRLTLLTSASAHYRIVVMG
Mytilus_californianus_XP_052084842  ----------MKSMKLLAVLPGNCIQSLSKNFSTGPASLQR-HFKVLVVG
Mytilus_californianus_XP_052084841  -------------MTSPKLLVVACRKSVTKSFSTSSASLGR-HYKLLIVG
SQOR_HUMAN                       MVPLVAVVSGPRAQLFACLLRLGTQQVGPLQLHTGASHAARNHYEVLVLG

SQRD-1_isoform_b                 GGAGGLGAASKFARKLPRGSVGIIEPREDHYYQPGFTLVGGGLMSLEANR
SQRD-1_isoform_c                 -------------------------------------------MSLEANR
SQRD-1_isoform_a                 GGAGGLGAASKFARKLPRGSVGIIEPREDHYYQPGFTLVGGGLMSLEANR
SQRD-1_isoform_f                 -------------------------------------------MSLEANR
SQRD-1_isoform_d                 --------------------------------------------------
SQRD-1_isoform_e                 --------------------------------------------------
Y9C9A.16_isoform_a               -------------------------------------------MTLDSNR
Y9C9A.16_isoform_b               GGAGGLGIASKFTRKLPSGSLGIIEPLEDHYYQPGFTLVGGGLMTLDSNR
Ascaris_suum_tr|F1L539           --------------------------------------------------
Ascaris_suum_tr|F1L970           GGTAGVAISNRFKNLVPKGQMAIIEPNKDHYYQGGFTLVAGGLKTVASCV
Mytilus_californianus_XP_052084842  AGAGGCSVANKFAHHVGGEKVGIIEPNDEHYYQPMWTLVGGGVKKLQDSV
Mytilus_californianus_XP_052084841  SGSGGCATASKFSKKLGAGQVGIIEPKDDHYYQPWFTLVGGGIKNVEDSG
SQOR_HUMAN                       GGSGGITMAARMKRKVGAENVAIVEPSERHFYQPIWTLVGAGAKQLSSSG

SQRD-1_isoform_b                 GKQKDLIPKNATWIQDKVQKFEPAKNSVKLRGGDEITYDYMVIAMGVQLR
SQRD-1_isoform_c                 GKQKDLIPKNATWIQDKVQKFEPAKNSVKLRGGDEITYDYMVIAMGVQLR
SQRD-1_isoform_a                 GKQKDLIPKNATWIQDKVQKFEPAKNSVKLRGGDEITYDYMVIAMGVQLR
SQRD-1_isoform_f                 GKQKDLIPKNATWIQDKVQKFEPAKNSVKLRGGDEITYDYMVIAMGVQLR
SQRD-1_isoform_d                 --------------------------------------------------
SQRD-1_isoform_e                 --------------------------------------------------
Y9C9A.16_isoform_a               KKQVNLIPKGATWIQDKVETFNPSQNTVVLRGGEEISYEYMVIAMGIHLR
Y9C9A.16_isoform_b               KKQVNLIPKGATWIQDKVETFNPSQNTVVLRGGEEISYEYMVIAMGIHLR
Ascaris_suum_tr|F1L539           --MASVLHKDNVWLKKSVAELRPKNNSVVLDDGSEIKYDLLIAALGLDIR
Ascaris_suum_tr|F1L970           KPTQALIHPNSVWIQKSIAELHPKNNSVILDDDSEIKYDILMVAPGLELR
Mytilus_californianus_XP_052084842  QKMSKLLPKKGTWMKTKAVAFDPEKCTVTTAEGEEVKYEYLIVSTGLQLN
Mytilus_californianus_XP_052084841  KKMSKVIPKKADWLKTTAEAFDPKNCTVTTATGEEVKYDFLVMACGLQLN
SQOR_HUMAN                       RPTASVIPSGVEWIKARVTELNPDKNCIHTDDDEKISYRYLIIALGIQLD

SQRD-1_isoform_b                 YDMIKGAKEALDT-PGVCSNYSPFYVEKHYKEAMNFKGGNALYTFPNTPI
SQRD-1_isoform_c                 YDMIKGAKEALDT-PGVCSNYSPFYVEKHYKEAMNFKGGNALYTFPNTPI
SQRD-1_isoform_a                 YDMIKGAKEALDT-PGVCSNYSPFYVEKHYKEAMNFKGGNALYTFPNTPI
SQRD-1_isoform_f                 YDMIKGAKEALDT-PGVCSNYSPFYVEKHYKEAMNFKGGNALYTFPNTPI
SQRD-1_isoform_d                 --------------------------------------------------
SQRD-1_isoform_e                 --------------------------------------------------
Y9C9A.16_isoform_a               FDMIKGAVEALET-PGVCSNYSPFHVQKHYQEVMNFQKGNAIYTFPNTPI
Y9C9A.16_isoform_b               FDMIKGAVEALET-PGVCSNYSPFHVQKHYQEVMNFQKGNAIYTFPNTPI
Ascaris_suum_tr|F1L539           FDMVEGLSEALHL-PGICSIYRHDLAEKTYRELQTFSQGNAVFTLPNTPI
Ascaris_suum_tr|F1L970           YDMVEGLPEALKL-KGVNSIYLPELAQKTDQELHSFKGGHALFTFPNTPI
Mytilus_californianus_XP_052084842  YNKIKGLPEAFDTDPTVCSNYDNYVQKTWPAIQNFKGGNAIFTFPNTPI
Mytilus_californianus_XP_052084841  YNLIKGLPEGFDKDPMICSNYDQYVQKTWPAIQKFKGGNAIFTLPNTPV
SQOR_HUMAN                       YEKIKGLPEGFAH-PKIGSNYSVKTVEKTWKALQDFKEGNAIFTFPNTPV

SQRD-1_isoform_b                 KCAGAPQKACYITDSILRQRGVRDQAHMIYATSLK---------------
SQRD-1_isoform_c                 KCAGAPQKACYITDSILRQRGVRDQAHMIYATSLK---------------
SQRD-1_isoform_a                 KCAGAPQKACYITDSILRQRGVRDQAHMIYATSLKRVRFLLCWDIINHGA
SQRD-1_isoform_f                 KCAGAPQKACYITDSILRQRGVRDQAHMIYATSLKRVRFLLCWDIINHGA
SQRD-1_isoform_d                 --------------------------------------------------
SQRD-1_isoform_e                 ---------------------------MIYATSLKRVRFLLCWDIINHGA
Y9C9A.16_isoform_a               KCAGAPQKACYITDSILRQRGVRDKAHMIYATSLS---------------
Y9C9A.16_isoform_b               KCAGAPQKACYITDSILRQRGVRDKAHMIYATSLS---------------
Ascaris_suum_tr|F1L539           KCSGAGQKICYLADEIFKKRGVRENIKLFYNTYLP---------------
Ascaris_suum_tr|F1L970           KCAGAPQKIMYISDEIFRQIGVRDKTKITYCTSLG---------------
Mytilus_californianus_XP_052084842  KCAGAPQKIMYLAEDWWRKNGMREQANIIFNSSLG---------------
Mytilus_californianus_XP_052084841  KCAGAPQKIMYLAEEAWQKSGVRDKATVMYNTSLS---------------
SQOR_HUMAN                       KCAGAPQKIMYLSEAYFRKTGKRSKANIIFNTSLG---------------
```

```
SQRD-1_isoform_b                   ----------------------------------RLFGIESYLKSLEKVA
SQRD-1_isoform_c                   ----------------------------------RLFGIESYLKSLEKVA
SQRD-1_isoform_a                   IGNQKSKKKIRKHQLVTIDCFDNNILKQYNNFFFHLFGIESYLKSLEKVA
SQRD-1_isoform_f                   IGNQKSKKKIRKHQLVTIDCFDNNILKQYNNFFFHLFGIESYLKSLEKVA
SQRD-1_isoform_d                   --------------------------MIYATSLKRLFGIESYLKSLEKVA
SQRD-1_isoform_e                   IGNQKSKKKIRKHQLVTIDCFDNNILKQYNNFFFHLFGIESYLKSLEKVA
Y9C9A.16_isoform_a                 ----------------------------------KIFGVDHYVKALEKVA
Y9C9A.16_isoform_b                 ----------------------------------KIFGVDHYVKALEKVA
Ascaris_suum_tr|F1L539             ----------------------------------DIFDVPKYAKTLNEIA
Ascaris_suum_tr|F1L970             ----------------------------------RVFGIKKYADALMKIV
Mytilus_californianus_XP_052084842 ----------------------------------VIFGVKKYANSLNKVI
Mytilus_californianus_XP_052084841 ----------------------------------VLFGVKKYAAGLHKVV
SQOR_HUMAN                         ----------------------------------AIFGVKKYADALQEII

SQRD-1_isoform_b                   RDKEIDVRTRRNLIEVNTNDRIATFELLDEEAKPTGKTEQIEYSLLHIGP
SQRD-1_isoform_c                   RDKEIDVRTRRNLIEVNTNDRIATFELLDEEAKPTGKTEQIEYSLLHIGP
SQRD-1_isoform_a                   RDKEIDVRTRRNLIEVNTNDRIATFELLDEEAKPTGKTEQIEYSLLHIGP
SQRD-1_isoform_f                   RDKEIDVRTRRNLIEVNTNDRIATFELLDEEAKPTGKTEQIEYSLLHIGP
SQRD-1_isoform_d                   RDKEIDVRTRRNLIEVNTNDRIATFELLDEEAKPTGKTEQIEYSLLHIGP
SQRD-1_isoform_e                   RDKEIDVRTRRNLIEVNTNDRIATFELLDEEAKPTGKTEQIEYSLLHIGP
Y9C9A.16_isoform_a                 QEKTIEVKTKRNLIEVVTNNKKAIFELLGDDSKPTGVTEEIEFSLLHISP
Y9C9A.16_isoform_b                 QEKTIEVKTKRNLIEVVTNNKKAIFELLGDDSKPTGVTEEIEFSLLHISP
Ascaris_suum_tr|F1L539             KQKHIDVSLRRSLKSIDPIKKEAVFDILTEQSKPSGKTVVQRYDLLHVAP
Ascaris_suum_tr|F1L970             ESKNIELNVRRNLIKVDPITRTATFEILDDDAVSTGKTVSFKYDFLHAAP
Mytilus_californianus_XP_052084842 ESRGMKVNYKRNLIEVRPKTREAIFQNLDSPT---GETETFKYEFLHVTP
Mytilus_californianus_XP_052084841 DSRDIKINYQRNLIEVNTDKREAIFQKLDSPS---GETETFKYDFMHITP
SQOR_HUMAN                         QERNLTVNYKKNLIEVRADKQEAVFENLDKP----GETQVISYEMLHVTP

SQRD-1_isoform_b                   PCSTPEALRNS-AFVDKTGFMDVDGGSLQSKKYPNVFGVGDCMNTPNAKT
SQRD-1_isoform_c                   PCSTPEALRNS-AFVDKTGFMDVDGGSLQSKKYPNVFGVGDCMNTPNAKT
SQRD-1_isoform_a                   PCSTPEALRNS-AFVDKTGFMDVDGGSLQSKKYPNVFGVGDCMNTPNAKT
SQRD-1_isoform_f                   PCSTPEALRNS-AFVDKTGFMDVDGGSLQSKKYPNVFGVGDCMNTPNAKT
SQRD-1_isoform_d                   PCSTPEALRNS-AFVDKTGFMDVDGGSLQSKKYPNVFGVGDCMNTPNAKT
SQRD-1_isoform_e                   PCSTPEALRNS-AFVDKTGFMDVDGGSLQSKKYPNVFGVGDCMNTPNAKT
Y9C9A.16_isoform_a                 PCTAPEPLRSS-HFADKTGFMDVDPQTLQSKNYPNVFGVGDCMNTPNAKT
Y9C9A.16_isoform_b                 PCTAPEPLRSS-HFADKTGFMDVDPQTLQSKNYPNVFGVGDCMNTPNAKT
Ascaris_suum_tr|F1L539             PCSPVKPIRECTELLNASGWLAADAATLQSTKFDNVFGIGDCLGTPNKKT
Ascaris_suum_tr|F1L970             PCSPVKPLRDCHELTDNKGWLDVDPDTLLSKKFDNVFGMGDCLNTSNAKT
Mytilus_californianus_XP_052084842 PMSTPDVVRDS-PLVDANGFVDVDKNTLQHKKYSNIFGIGDCTNAPTSKT
Mytilus_californianus_XP_052084841 PMSTPDPVRKS-CLVDEAGYVDVDQGTLQHKKYPNIFAIGDCFNAPTSKT
SQOR_HUMAN                         PMSPPDVLKTS-PVADAAGWVDVDKETLQHRRYPNVFGIGDCTNLPTSKT

SQRD-1_isoform_b                   AAAVSSHLKTIEKNLTQVMQGNRPCMQYDGYASCPLVVSTNRVILAEFGP
SQRD-1_isoform_c                   AAAVSSHLKTIEKNLTQVMQGNRPCMQYDGYASCPLVVSTNRVILAEFGP
SQRD-1_isoform_a                   AAAVSSHLKTIEKNLTQVMQGNRPCMQYDGYASCPLVVSTNRVILAEFGP
SQRD-1_isoform_f                   AAAVSSHLKTIEKNLTQVMQGNRPCMQYDGYASCPLVVSTNRVILAEFGP
SQRD-1_isoform_d                   AAAVSSHLKTIEKNLTQVMQGNRPCMQYDGYASCPLVVSTNRVILAEFGP
SQRD-1_isoform_e                   AAAVSSHLKTIEKNLTQVMQGNRPCMQYDGYASCPLVVSTNRVILAEFGP
Y9C9A.16_isoform_a                 AASVSTHLKTVDTNLTQVMQGARPFMKYDGYASCPLVVSTNRVILAEFSP
Y9C9A.16_isoform_b                 AASVSTHLKTVDTNLTQVMQGARPFMKYDGYASCPLVVSTNRVILAEFSP
Ascaris_suum_tr|F1L539             SAAVFSQLRALDKNIPAFLSGKKLEGQYNGYSSCPLIVSFNRVVLAEFTP
Ascaris_suum_tr|F1L970             GAAISSQMRVMSKNLPALLHGKPLTGKYDGYASCPLMVAMNKVILAEFNS
Mytilus_californianus_XP_052084842 AAAAAAQCGILETNLKAVMADKPMTAQYDGYTSCPLITARGKCILAEFDF
Mytilus_californianus_XP_052084841 AAAAASQSGYLEKNLTAVMNGKPAYVMYDGYTSCPLITGREKCIMAEFNY
SQOR_HUMAN                         AAAVAAQSGILDRTISVIMKNQTPTKKYDGYTSCPLVTGYNRVILAEFDY

SQRD-1_isoform_b                   RG-AMETTPFDQSKPTYWAYLMKRYFMPALYWNGLIKGYWNGPATLRNCT
SQRD-1_isoform_c                   RG-AMETTPFDQSKPTYWAYLMKRYFMPALYWNGLIKGYWNGPATLRNCT
SQRD-1_isoform_a                   RG-AMETTPFDQSKPTYWAYLMKRYFMPALYWNGLIKGYWNGPATLRNCT
SQRD-1_isoform_f                   RG-AMETTPFDQSKPTYWAYLMKRYFMPALYWNGLIKGYWNGPATLRNCT
SQRD-1_isoform_d                   RG-AMETTPFDQSKPTYWAYLMKRYFMPALYWNGLIKGYWNGPATLRNCT
SQRD-1_isoform_e                   RG-AMETTPFDQSKPTYWAYLMKRYFMPALYWNGLIKGYWNGPATLRNCT
Y9C9A.16_isoform_a                 NE-ILETTPLNQSKPSYWAFLMKRYILPVLYWKGLIKGHWNGPSTIRNFS
Y9C9A.16_isoform_b                 NE-ILETTPLNQSKPSYWAFLMKRYILPVLYWKGLIKGHWNGPSTIRNFS
Ascaris_suum_tr|F1L539             KG-PLETLPYDQGRPLYTAYLLKRYVLPTMYWTIGINGHWLGPATVRKIL
Ascaris_suum_tr|F1L970             DG-PLETLPINGAKPRYSSYLLKRYFLPFLYWNFLIKGLWLGP-------
Mytilus_californianus_XP_052084842 DGNPLETFPINQGKERMTMYHMKKDVMPQIYWHMMMSGHWNGPGVYRKLM
Mytilus_californianus_XP_052084841 DSLPLETFPIDQGKERRSMYHVKKDIIPMIYWNMMLKGYWNGPAAYRKLM
SQOR_HUMAN                         KAEPLETFPFDQSKERLSMYLMKADLMPFLYWNMMLRGYWGGPAFLRKLF

SQRD-1_isoform_b                   RLVKSK--------------------------------------------
SQRD-1_isoform_c                   RLVKSK--------------------------------------------
SQRD-1_isoform_a                   RLVKSK--------------------------------------------
SQRD-1_isoform_f                   RLVKSK--------------------------------------------
```

```
SQRD-1_isoform_d                    RLVKSK--------------------------------------------
SQRD-1_isoform_e                    RLVKSK--------------------------------------------
Y9C9A.16_isoform_a                  RLLKTK--------------------------------------------
Y9C9A.16_isoform_b                  RLLKTK--------------------------------------------
Ascaris_suum_tr|F1L539              HLRRSS--------------------------------------------
Ascaris_suum_tr|F1L970              --------------------------------------------------
Mytilus_californianus_XP_052084842  HLGMDGPEKQAASA------------------------------------
Mytilus_californianus_XP_052084841  HFGTDGPEKEKDSY------------------------------------
SQOR_HUMAN                          HLGMS---------------------------------------------
```

**Figure S1**. Alignment of sulfide:quinone oxidoreductases from *C. elegans* and other animal lineages that possess two SQRD genes with the human protein. *C. elegans* possess 5 isoforms derived from the *sqrd-1* gene and two derived from the Y9C9A.16 *(sqrd-2*) gene.  Rossmann domains are highlithed in green. The sulfide contact residues are highlighted in yellow, and the key residues that form hydrogen bond with quinones are highlighted in cyan.

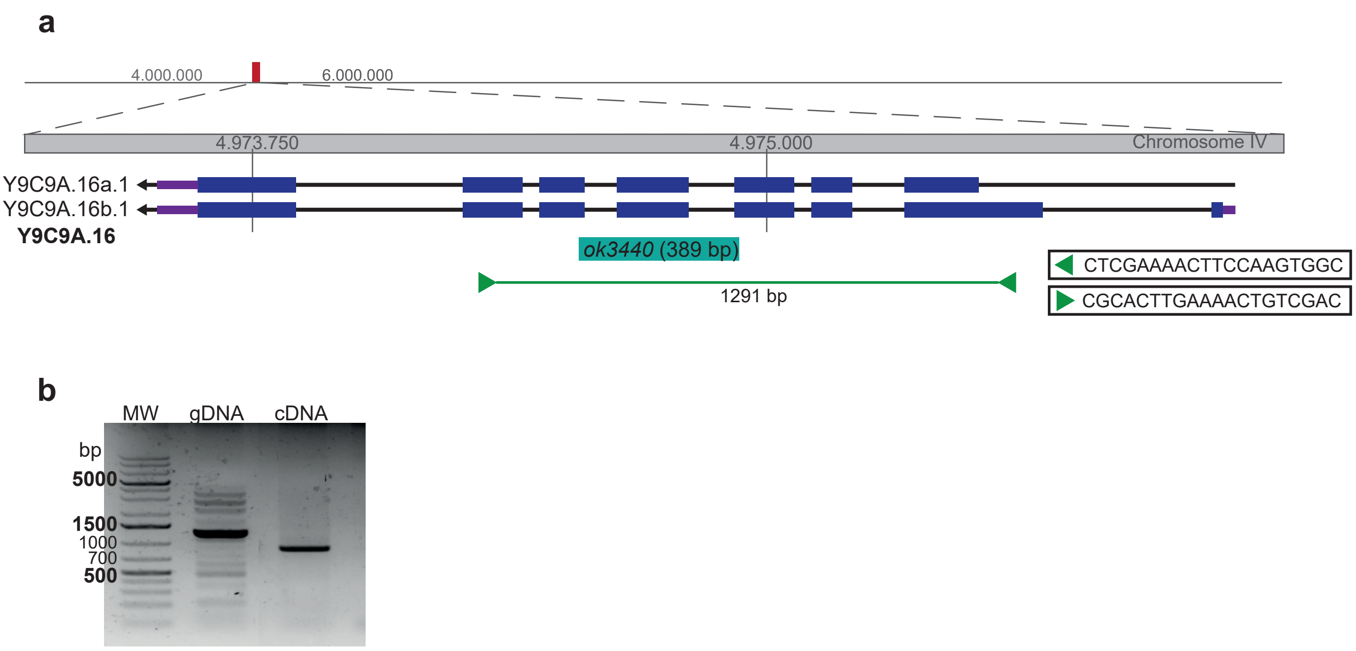


**Figure S2:** *C. elegans Y9C9A.16* (*sqrd-2*) gene model and mRNA PCR amplification product. **a**) Gene model of *Y9C9A.16* obtained from the wormbase.org. Y9C9A.16a.1 and Y9C9A.16b.1 are two predicted isoforms. *ok3440* is the mutant allele used in this work (389 bp deletion size) and the bar indicate its location within the gene. Green arrowheads and line represent the oligo primers used for PCR and the size and gene region amplified, respectively. **b**) PCR products obtained with oligo primers listed in **a**) using genomic DNA (gDNA) or cDNA as template.

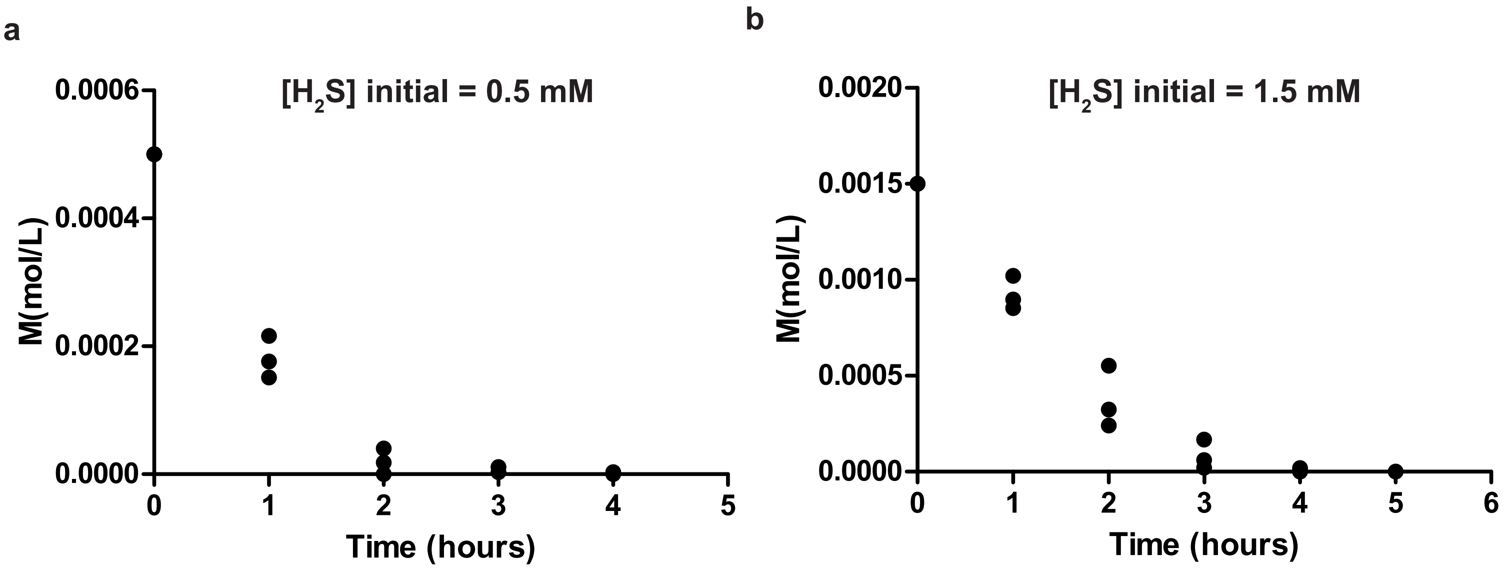


**Figure S3:** $H_2S$ concentration in solution decay over the time. $H_2S$ concentration in solution was determined using the colorimetric assay described by Siegel, L. M., 1965. Concentration (mol/L) at different time points using 1.5 mM (**a**) or 0.5 mM (**b**) of $Na_2S$ as initial concentration were determined in three different experiments.

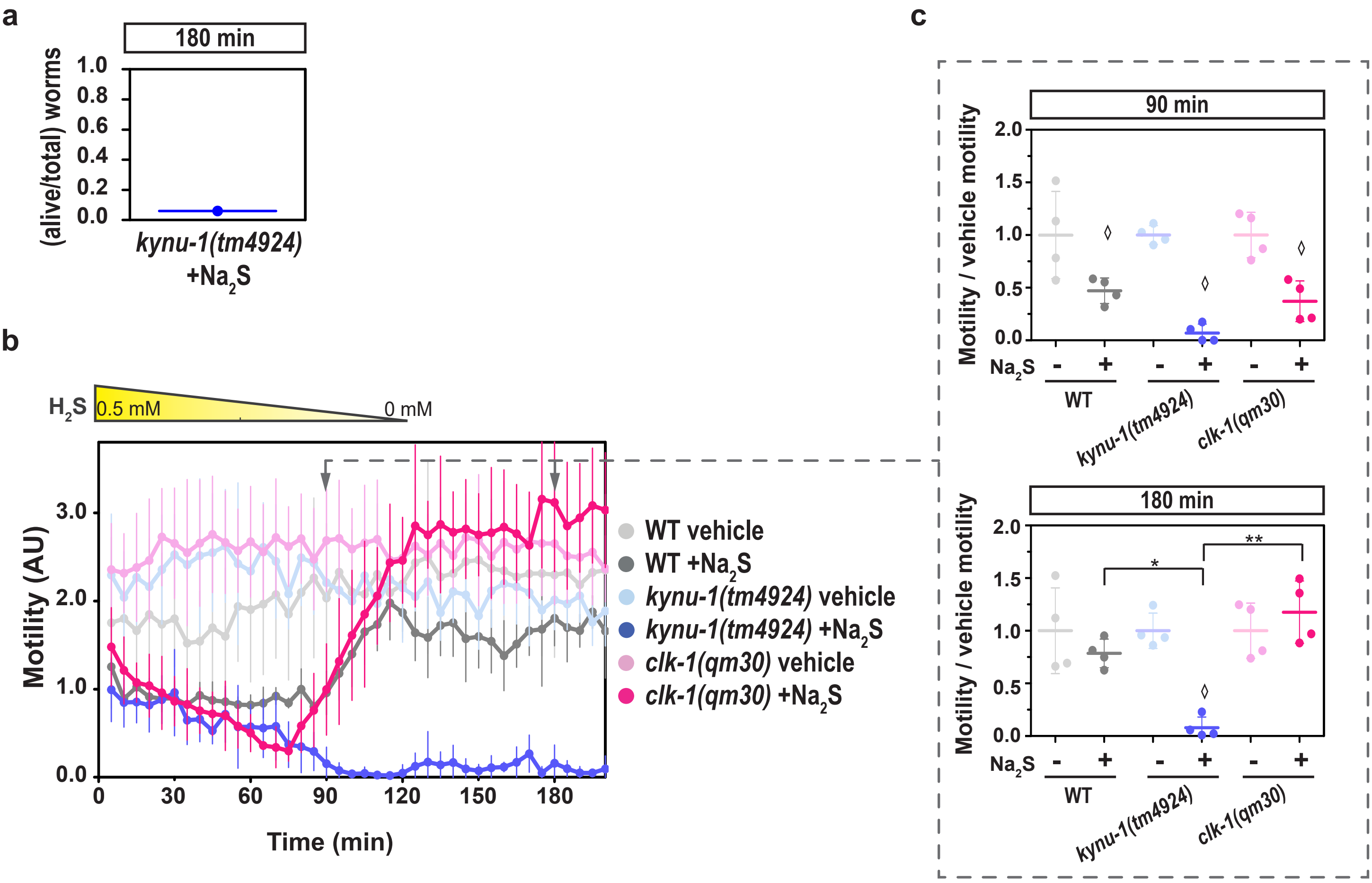


**Figure S4:** The rhodoquinone-deficient mutant strain (*kynu-1(tm4924)*) does not recover from sulfide challenge in contrast to the ubiquinone-deficient mutant strain (*clk-1(qm30)*). **a**) The graph indicates the survival (ratio: number of alive worms/total number of worms) of *kynu-1(tm4924)* mutant animals after 3 hours in $Na_2S$ (initial concentration=1.5 mM), n=1 replicate, 50 animals. **b**) The motility parameter refers to the movement of a population of individuals in liquid media and was measured using the infrared tracking device WMicrotracker. Motility of *kynu-1(tm4924)* and *clk-1(qm30)* mutants and wild-type (WT) strains in the presence (+$Na_2S$) or absence (vehicle) of a sodium sulfide ($Na_2S$) solution. Each point indicates the motility average of 4 wells (relative to the habituation, see methods) measured every 5 minutes for 200 minutes. The graph corresponds to a representative experiment with 4 wells per condition per strain (approximately 80 worms per well). Error bars indicate the standard deviation. At least three biological replicates were performed for each worm strain. The yellow triangle represents the decrease of the sulfide concentration in solution from 0.5 mM. Arrows represent the two time points shown in part **c**). AU: arbitrary units. **c**) For each strain *(kynu-1(tm4924)*, *clk-1(qm30)* and WT), the motility in the presence (+) or absence (-) of sulfide ($Na_2S$) was normalized to the mean of the motility in the absence of $Na_2S$. Each data point represents the normalized motility of each one of the 4 wells for the two conditions (with or without $Na_2S$). Different graphs correspond to the time point of 90 and 180 minutes of incubation. Welch test (90 min p=1,98E-7 and 180 min p=5,2E-7) were performed, followed by Tukey´s pairwise test. Asterisks indicate statistical differences (180min: *p=3,9E-3 and **p=3,2E-4). Diamonds indicate statistical differences for each strain between sulfide and their control without sulfide. The p values obtained with statistical tests are reported in Supporting information.

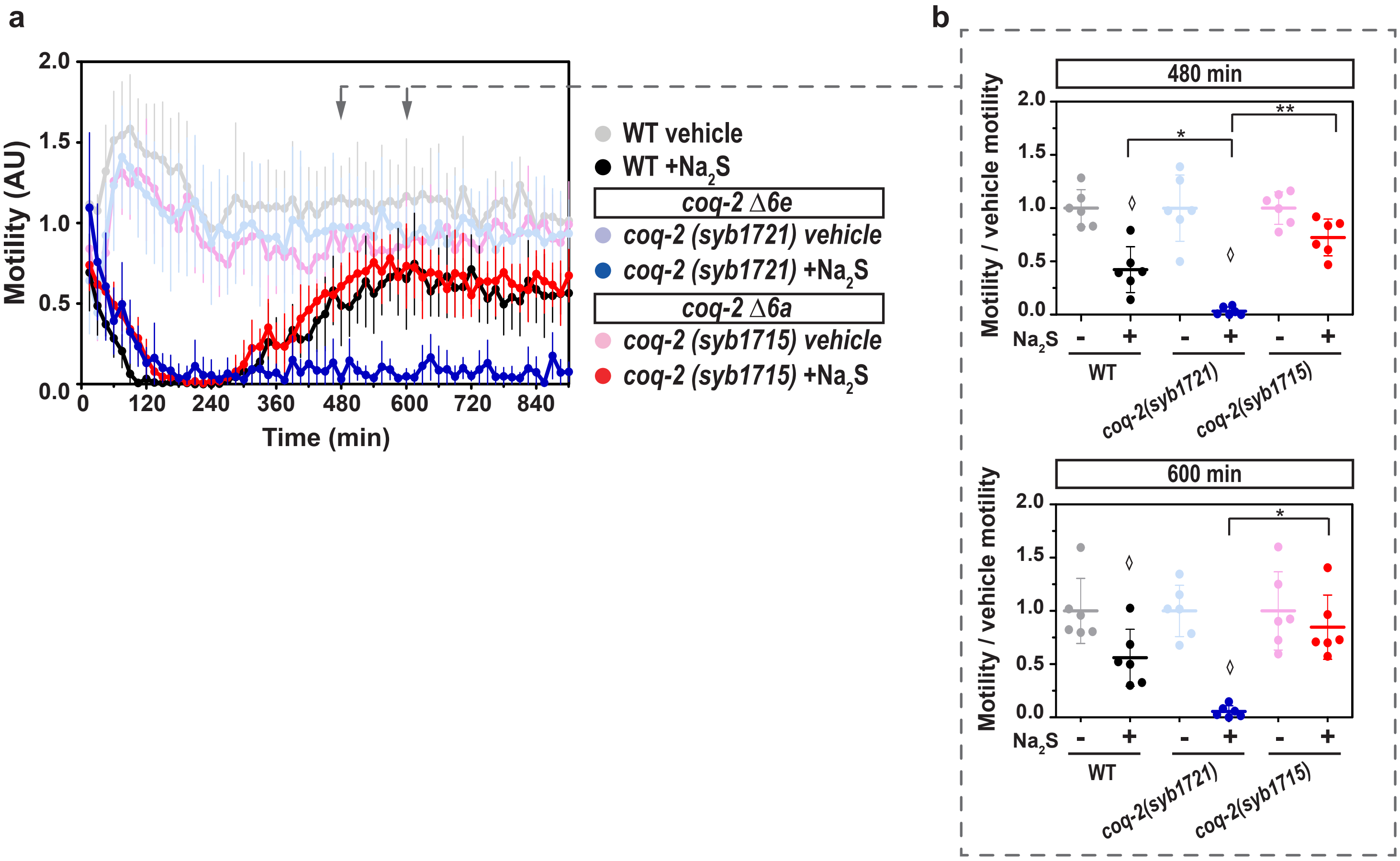


**Figure S5:** The rhodoquinone-deficient mutant strain (*coq-2(syb1721)*, *(coq-2Δ6e)*) does not recover from sulfide challenge in contrast to the ubiquinone-deficient mutant strain (*coq-2(syb1715)*, *(coq-2Δ 6a)*). The motility parameter refers to the movement of a population of individuals in liquid media and was measured using the infrared tracking device *WMicrotracker.* **a**) Motility of *coq-2(syb1721)*, *coq-2(syb1715)* mutant and wild-type (WT) strains in the presence (+$Na_2S$) or absence (vehicle) of a sodium sulfide ($Na_2S$) solution. The vehicle used was destilled water. Each point indicates the motility average of 6 wells (relative to the habituation, see methods) measured every 15 minutes for 900 minutes. The graph corresponds to a representative experiment with 6 wells per condition per strain (approximately 80 worms per well). Error bars indicate the standard deviation. At least three biological replicates were performed for each worm strain. Arrows represent the two time points shown in part **b**). AU: arbitrary units. **b**) For each strain (*coq-2(syb1721)*, *coq-2(syb1715)* and WT), the motility in the presence (+) or absence (-) of sulfide ($Na_2S$) was normalized to the mean of the motility in the absence of $Na_2S$. Each data point represents the normalized motility of each one of the 6 wells for the two conditions (with or without $Na_2S$). Different graphs correspond to the time point of 480 and 600 minutes of incubation. ANOVA test (480 min p=7,05E-10) followed by Tukey´s pairwise test and Kruskal-Wallis test (600 min p=1,0E-3) followed by Dunn´s pairwise test were performed. Asterisks indicate statistical differences (480 min: *p=0.018 and **p=1.3E-5; 600 min: *p=6.7E-3). Diamonds indicate statistical differences for each strain between sulfide and their control without sulfide. The p values obtained with statistical tests are reported in Supporting information.

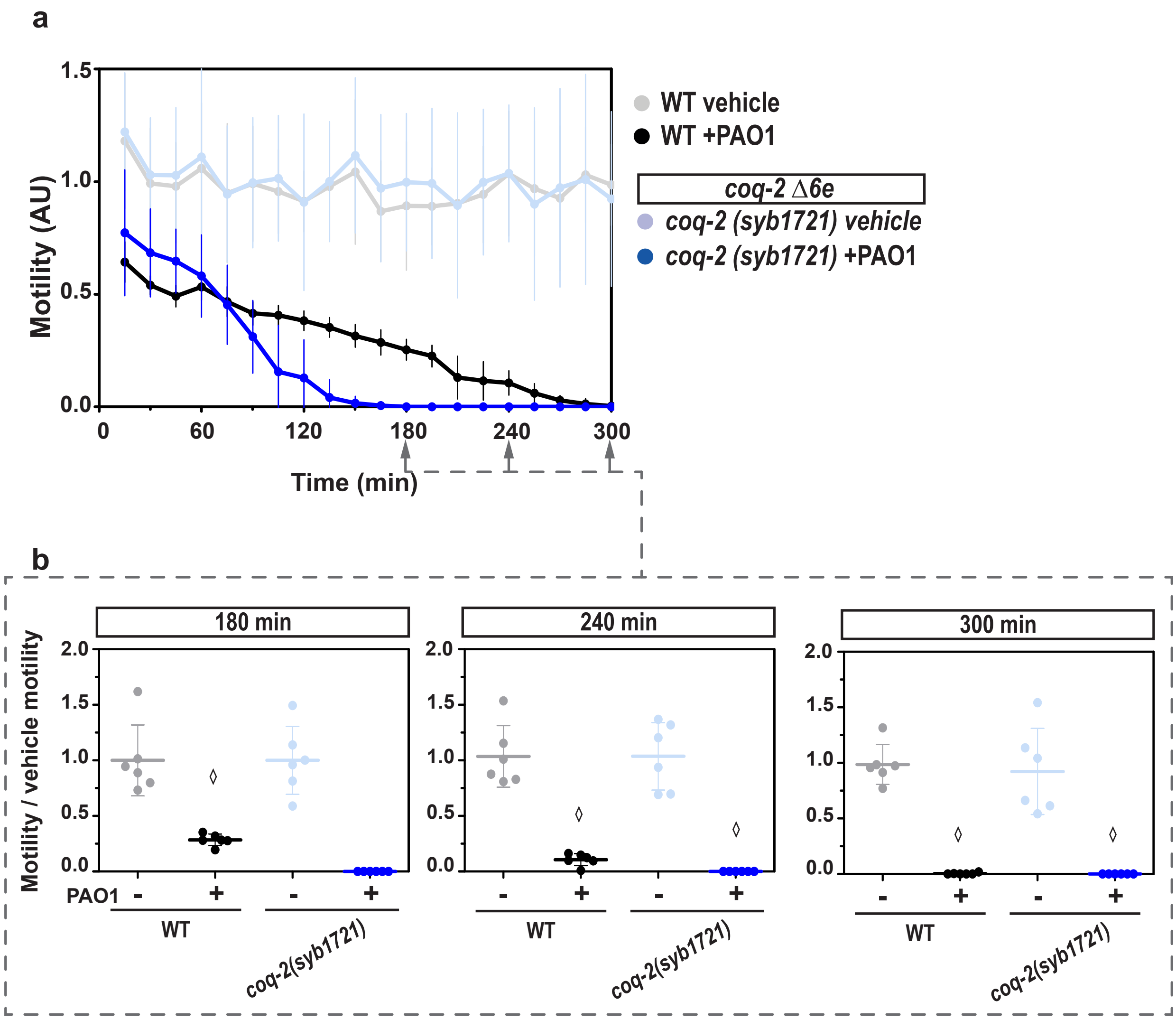


**Figure S6:** The rhodoquinone-deficient mutant strain *coq-2(syb1721)* (*coq-2Δ 6e*) is more sensitive to *Pseudomonas aeruginosa* PAO1 strain than the wild-type strain. The motility parameter refers to the movement of a population of individuals in liquid media and was measured using the infrared tracking device *WMicrotracker.* **a**) Motility of *coq-2(syb1721)* and wild-type (WT) strains in the presence (+PAO1) or absence (vehicle) of a liquid culture of *P. aeruginosa* PAO1. Each point indicates the motility average of 6 wells (relative to the habituation, see methods) measured every 15 minutes for 300 minutes. The graph corresponds to a representative experiment with 6 wells per condition per strain (approximately 80 worms per well). Error bars indicate the standard deviation. At least three biological replicates were performed for each worm strain. Arrows represent the time points shown in part **b**). AU: arbitrary units. **b**) For each strain (*coq-2(syb1721)* and WT), the motility in the presence (+) or absence (-) of P. *aeruginosa* PAO1 was normalized to the mean of the motility in the absence of the bacteria. Each data point represents the normalized motility of each one of the 6 wells for the two conditions (with or without bacteria). Different graphs correspond to the time point of 180, 240 and 300 minutes of incubation. Kruskal-Wallis test (180 min p=1.9E-4, 240 min p=1.9E-4 and 300 min p=2.9E-4) were performed, followed by Dunn´s post hoc test. Diamonds indicate statistical differences for each strain between the two conditions (bacteria and vehicle). All the p values obtained with different statistic tests are reported in Supporting information.

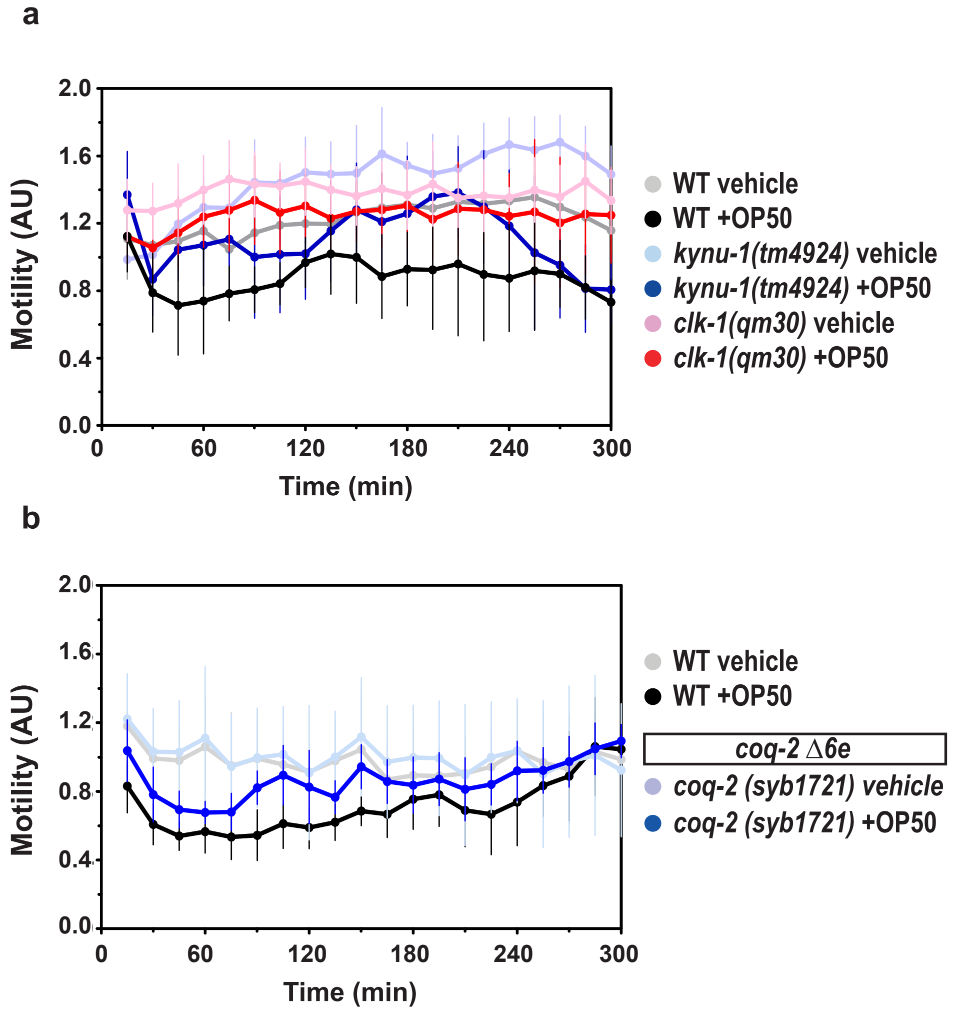


**Figure S7:** Liquid culture of the food bacterium *Escherichia coli* OP50 does not affect differentially the motility of the mutant strains *kynu-1(tm4924)*, *clk-1(qm30)*, *coq-2(syb1721)(coq-2Δ 6e)*, in comparison with the wild-type strain. The motility parameter refers to the movement of a population of individuals in liquid media and was measured using the infrared tracking device *WMicrotracker.* **a**) Motility of *kynu-1(tm4924)* and *clk-1(qm30)* and wild-type (WT) strains in the presence (+OP50) or absence (vehicle) of a liquid culture of *E. coli* OP50. Each point indicates the motility average of 6 wells (relative to the habituation, see methods) measured every 15 minutes for 300 minutes. The graph corresponds to a representative experiment with 6 wells per condition per strain (approximately 80 worms per well). Error bars indicate the standard deviation. At least three biological replicates were performed for each worm strain. **b**) The same as in part **a**) but with the strains *coq-2(syb1721)* and wild-type (WT).
